# The 3D architecture and molecular foundations of *de novo* centriole assembly *via* bicentrioles

**DOI:** 10.1101/2020.12.21.423647

**Authors:** Sónia Gomes Pereira, Ana Laura Sousa, Catarina Nabais, Tiago Paixão, Alexander. J. Holmes, Martin Schorb, Gohta Goshima, Erin M. Tranfield, Jörg D. Becker, Mónica Bettencourt-Dias

## Abstract

Centrioles are structurally conserved organelles, composing both centrosomes and cilia. In animal cycling cells, centrioles often form through a highly characterized process termed canonical duplication. However, a large diversity of eukaryotes form centrioles *de novo* through uncharacterized pathways. This unexplored diversity is key to understanding centriole assembly mechanisms and how they evolved to assist specific cellular functions. Here, combining electron microscopy and tomography, we show that during spermatogenesis of the moss *Physcomitrium patens*, centrioles are born as a co-axially oriented centriole pair united by a cartwheel. We observe that microtubules emanate from those bicentrioles, which localize to the spindle poles during cell division. Thereafter, each bicentriole breaks apart, and the two resulting sister centrioles mature asymmetrically, elongating specific microtubule triplets and a naked cartwheel. Subsequently, two cilia are assembled which are capable of beating asynchronously. We further show that conserved cartwheel and centriole wall components, SAS6, BLD10 and POC1 are expressed during spermatogenesis and are required for this *de novo* biogenesis pathway. Our work supports a scenario where centriole biogenesis is more diverse than previously thought and that conserved molecular modules underlie diversification of this essential pathway.

## Introduction

Centrioles are microtubule (MT)-based structures that compose centrosomes and cilia. The centrosome is the main microtubule organizing center (MTOC) in most animal cells, regulating intracellular transport, spindle pole formation and cell migration. Centrioles, then called basal bodies, can also anchor to the cell membrane and template cilia growth, with important roles in cell signaling and motility (reviewed in ref. Joukov and De Nicolo, 2019). Centriole biogenesis is tightly regulated in space, time and number. Failure in regulating this process can lead to diseases, such as cancer (Levine *et al*., 2017; Lopes *et al*., 2018; Marteil *et al*., 2018) and ciliopathies (Khan *et al*., 2014; Shaheen *et al*., 2012).

Centrioles are widespread across eukaryotes, being assembled by numerous pathways. The most prevalent and well-characterized of such pathways is canonical centriole duplication, a process by which one daughter centriole is assembled orthogonally to a pre-existing one (its mother) in a cell-cycle dependent manner. Accordingly, mature centrioles impose a numerical, spatial and temporal regulation on centriole duplication, which has been extensively characterized (reviewed in ref. Nigg and Holland, 2018). However, centrioles can also assemble *de novo* without pre-existing ones, raising the question of how is such a process regulated (Khodjakov *et al*., 2002; Miki-Noumura, 1977; Peel *et al*., 2007). This significantly less studied mechanism occurs for instance in vertebrate cells that form multiple cilia (Mercey *et al*., 2019; Sorokin, 1968). Moreover, it is clear from older electron microscopy studies that *de novo* centriole biogenesis is spread across eukaryotes, underlying the diversification of essential cellular functions such as stress evasion, spermatogenesis and embryo development (reviewed in ref. Nabais *et al*., 2018). The regulation, structures and molecules underpinning *de novo centriole* biogenesis remain vastly overlooked, in part due to the lack of amenable model systems and tools required to tackle this problem. This knowledge is key to shedding light on the evolutionary history and general principles governing centriole assembly.

Early land plants (such as bryophytes), ferns and some gymnosperms (*Ginkgo* and cycads) are critical ecological players, which reproduce through means of multiciliated motile sperm cells (reviewed in ref. Renzaglia and Garbary, 2001). In these plant species, sperm cells are the only centriole-containing cells in the entire organism. Therefore, during spermatogenesis either two (*e.g.* bryophytes) (Moser and Kreitner, 1970; Robbins, 1984) or many (*e.g. Ginkgo biloba*) (Gifford and Larson, 1980) centrioles arise *de novo*, through mechanisms that have remained poorly understood despite the importance of understanding their reproductive process.

In this work, we have used the moss *Physcomitrium patens (P. patens*, previously known as *Physcomitrella patens*) to investigate naturally occurring *de novo* centriole biogenesis in early land plants. Traditionally used as a model to study plant evolution (Prigge and Bezanilla, 2010; Rensing *et al*., 2008), the bryophyte *P. patens* reproduces via motile biciliated sperm cells, which develop inside specialized male organs called antheridia (Landberg *et al*., 2013). In addition to its simple anatomy, a haploid-dominant life-cycle and availability of genetic engineering tools make *P. patens* an attractive model for cell biology studies (Rensing *et al*., 2020). Moreover, early electron microscopy studies had described the formation of intriguing structures, two linked centrioles, at the origin of the locomotory apparatus of two other early land plant species (Moser and Kreitner, 1970; Robbins, 1984), raising important questions about the architecture, molecular composition and function of such structure.

Here, armed with modern molecular, light and electron microscopy tools, we explore the cell biology of centriole and cilia assembly during spermatogenesis in *P. patens*. High end 3D electron tomography uncovered the bicentriole-mediated pathway for centriole biogenesis with unprecedented detail, revealing unknown and surprising structures. Unexpectedly, this pathway results in distinctive centrioles that bear long portions of MT-deprived (“naked”) cartwheels and MT triplets of different lengths. These asymmetric centrioles form cilia that frequently display an asynchronous beating pattern. The diversity of the observed structures begged the question of whether the same molecular players known to other organisms are involved in bicentriole assembly and maturation. We investigated the role of the conserved molecular players SAS6, BLD10 and POC1 during centriole assembly and maturation, showing their requirement in assembling the diverse centriole structures. Our work suggests that an evolutionary conserved centriole assembly module is required for the biogenesis of diverse centriole structures, underlying critical cellular and organismal functions such as motility and fertility.

## Results

### The bicentriole-mediated pathway for *de novo* centriole biogenesis

As a first approach to investigate centriole assembly in *P. patens* we used transmission electron microscopy (TEM). The centriole is known for its 9-fold symmetry, complex ultrastructure and small size (reviewed in LeGuennec *et al*., 2020). Centriole biogenesis has been studied in different organisms using several electron microscopy (EM) techniques, as these offer a unique opportunity to study small cellular structures with high level of detail and are not constrained by antibody availability, nor molecular conservation. *P. patens’* spermatogenesis occurs asynchronously amongst the several antheridia from the same individual, with the first sperm cells maturing within 15 days after induction (DAI) of sexual reproduction (Fig. 1A). We used TEM imaging of samples fixed at successive DAI to define the order of events that characterize *de novo* centriole biogenesis during *P. patens’* spermatogenesis (Fig. 1B), building an overview of the pathway (Fig. 1C) that guided us in further analyses.

**Figure 1.**
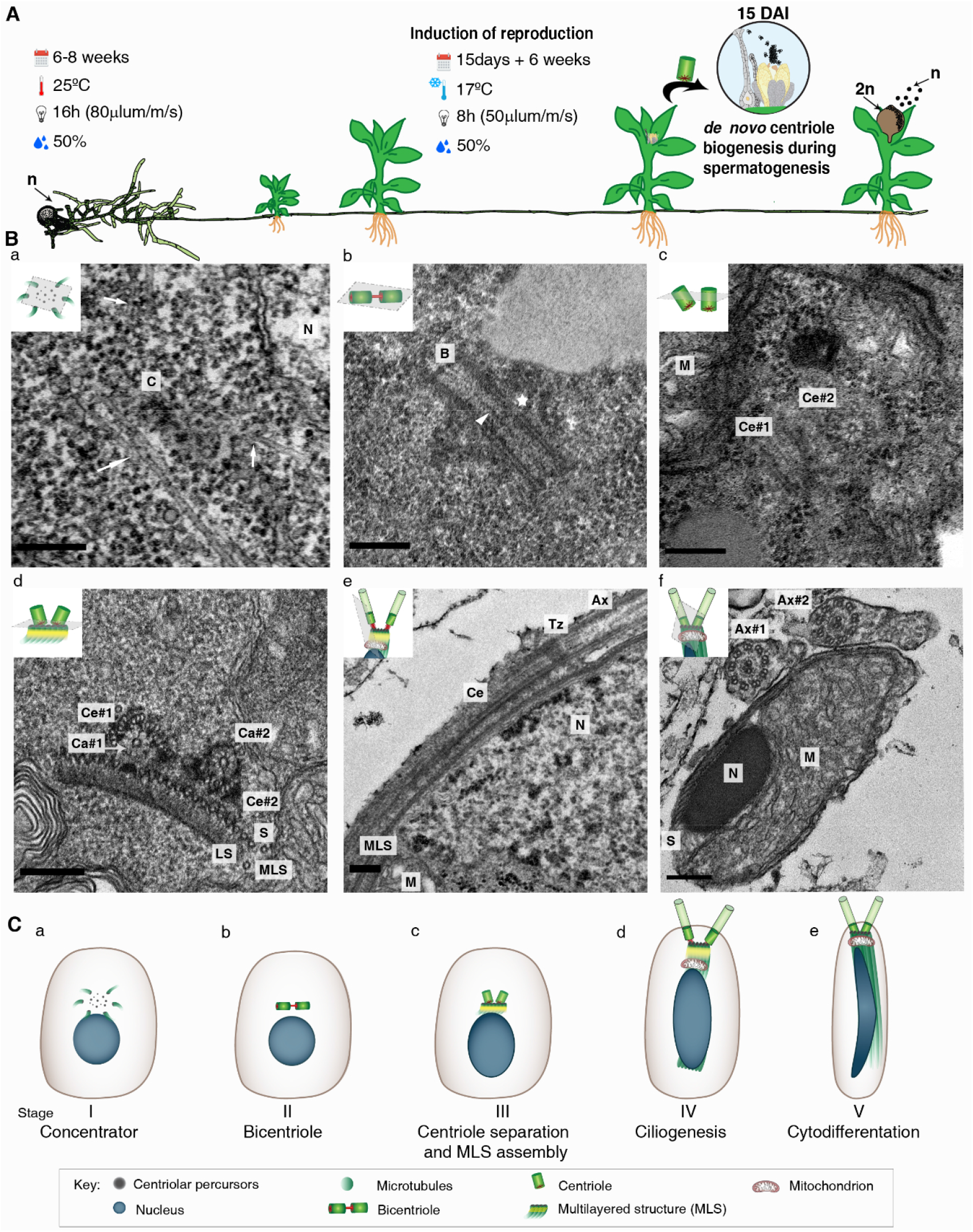
The bicentriole-mediated pathway for *de novo* centriole biogenesis. **A,** The haploid dominant life-cycle of *P. patens*. It is only during spermatogenesis that centrioles arise *de novo*. After a period of 6 to 8 weeks of vegetative growth, sexual reproduction is induced and 15 Days After Induction (15DAI) the first biflagellated sperm cells are released. Fertilization gives rise to a sporophyte, which represents the only diploid (2n) stage of the life-cycle. Meiosis occurs in this sporophyte culminating in the release of haploid (n) spores. **B,** TEM representative images of the bicentriole-mediated centriole assembly. **Ba,** Concentrator (C) with microtubules emanating from it (arrows). **Bb,** Bicentriole (B), arrowhead highlighting the continuous cartwheel and the star the discontinuity of the centriolar wall. **Bc,** Individualized centrioles organized side by side. **Bd**, Sister centrioles docked to the multilayered structure (MLS). **Be,** Longitudinal view of a centriole docked to the cellular membrane and templating the growth of the ciliary axoneme. **Bf**, Cell cytodifferentiation yields an elongated mitochondrion and condensed nucleus. C – concentrator; N – nucleus; B – bicentriole; M - mitochondrion; Ce#1 and Ce#2 – centrioles numbered 1 and 2 (numbering is arbitrary); Ca#1 and Ca#2 – cartwheel from centriole #1 and #2, respectively; MLS – multilayered structure, composed of the spline (S) and lamellar strip (LS); Tz: transition zone; Ax: ciliary axoneme (Ax#1 and Ax#2 – axonemes number 1 and 2, arbitrary numbering). Scale-bars = 200nm. For sample size, please refer to Table S1. **C,** Schematic of the bicentriole-mediated pathway in *P. patens*. **Ca,** Concentrator – stage I. **Cb,** Bicentriole – stage II. **Cc,** Separated centrioles and MLS assembly – stage III. **Cd,** Centrioles docked to the membrane and ciliogenesis – stage IV. **Ce,** Sperm cell cytodifferentiation – stage V.

At early stages of spermatogenesis (10-12 DAI) we failed to detect any centriole-like structures within sperm cells. However, we observed near the nucleus a cloud of electron-dense unstructured material with MTs emanating from it (Fig. 1Ba). This electron dense material, may act as an MTOC and precede centriole biogenesis, perhaps concentrating its precursors, and therefore we named it “concentrator” (Fig. 1Ca). From 13 DAI onwards, we started to observe bicentriole-like structures (Fig. 1Bb, 1Cb). These are composed of a continuous cartwheel (Fig. 1Bb arrowhead), which is surrounded by a MT wall that shows a small discontinuity between two co-axially oriented centrioles (Fig. 1Bb star).

As spermatogenesis proceeds, two individual centrioles were observed (Fig. 1Bc) per sperm cell, likely resulting from splitting of the bicentriole. Below the individualized centrioles, a multilayered structure (MLS) was recognizable. The MLS is composed of a spline of parallel singlet MTs on top of electron dense protein plates called lamellar strip (LS) (Carothers and Duckett, 1980) (Fig. 1Bd, 1Cc). The complex consisting of the two centrioles and the MLS migrated towards the cell surface, after which the centrioles docked to the membrane (Fig. 1Be). Both centrioles became basal bodies, assembling two axonemes with the characteristic 9+2 MT arrangement of motile cilia (Fig. 1Bf, 1Cd).

At this point, the sperm cells’ locomotory apparatus was composed of two centrioles and cilia, as well as the plant-specific MLS (Fig. 1Cd). The final step of cytodifferentiation (14-15DAI) involved cytosolic volume reduction, and mitochondria fusion, followed by chromatin condensation and nuclear elongation (Fig. 1Bf, 1Ce). Interestingly, the lamellar strip component of the MLS disappeared in the final stage of development (Fig. 1Bf), which is similar to what is found in other plant species, such as *Marchantia polymorpha* and *Phaeoceros laevis* (Carothers and Duckett, 1980).

Our analysis showed several intriguing features of *P. patens* centrioles, namely the frequent observation of incomplete centriolar walls (Fig. 1Bd) and long cartwheel structures with no attached MTs (Fig. S1A). These may reflect unknown steps of biogenesis and/or maturation that are unfortunately difficult to capture or visualize as we only have 2D information. In order to further understand the organization of *P. patens* locomotory apparatus, we proceeded to perform 3D electron tomography (ET) throughout different maturation stages (Fig. 2–3).

**Figure 2.**
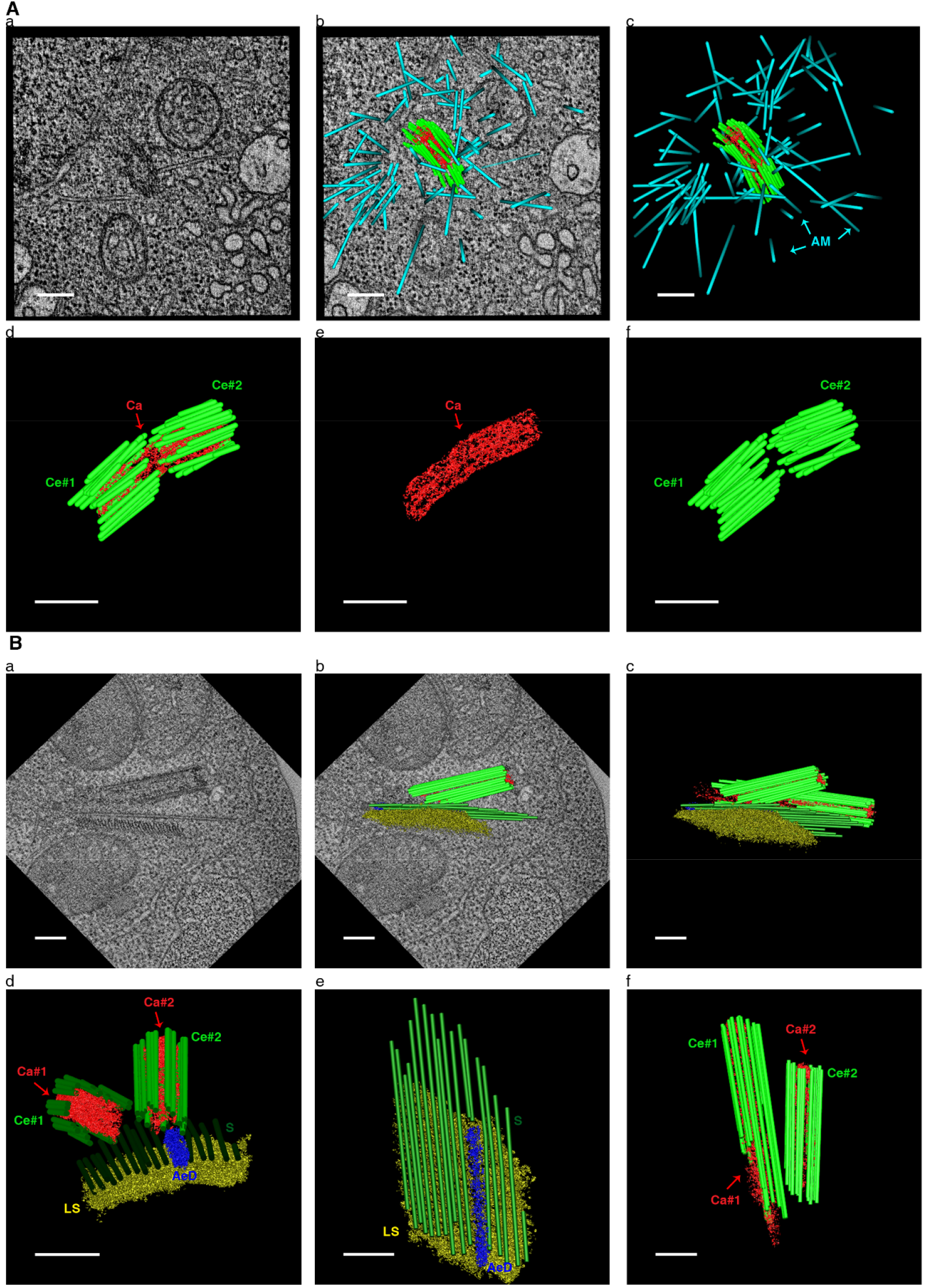
3D analysis of a bicentriole and MLS-anchored centrioles. **A,** 3D structure of a bicentriole (see movie 1). **Aa,** Electron tomogram snapshot. **Ab,** Superimposition of the tomogram with its segmentation. **Ac,** 3D model of a bicentriole, with several astral microtubules emanating from the structure. **Ad,** Rotated 3D model of the bicentriole, composed of two centriolar units connected by their inner continuous cartwheel. **Ae**, Continuous cartwheel that connects both centrioles. **Af,** The walls of the two centrioles of the same bicentriole are discontinuous. **B,** 3D architecture of individualized sister centrioles anchored to the MLS (see movie 2). **Ba,** Snapshot of an electron tomogram showing a longitudinal view of one centriole anchored to the MLS. Striation is visible on the lamellar strip. **Bb,** Electron tomogram and its 3D model overlapping. **Bc,**3D model of centrioles anchored to the MLS. **Bd,** Rotated view of the 3D model represented in Bc, highlighting the existence of an amorphous electron density (AeD) between the LS and one of the centrioles (Ce#2). **Be,** Top view of the MLS structure. The AeD is located in a gap of the spline’s microtubules and directly above the LS. **Bf,** Top view of the two individualized centrioles. AM (cyan) – astral microtubules; Ce #1 and #2 (light green) – wall of centrioles numbered 1 and 2 (arbitrary numbering); Ca #1 and #2 (red) – cartwheel of centrioles number #1 or #2, respectively; LS (yellow) – lamellar strip of the MLS; S (dark green) – spline of the MLS; AeD (blue) – amorphous electron density. Scale-bars = 200nm.

**Figure 3.**
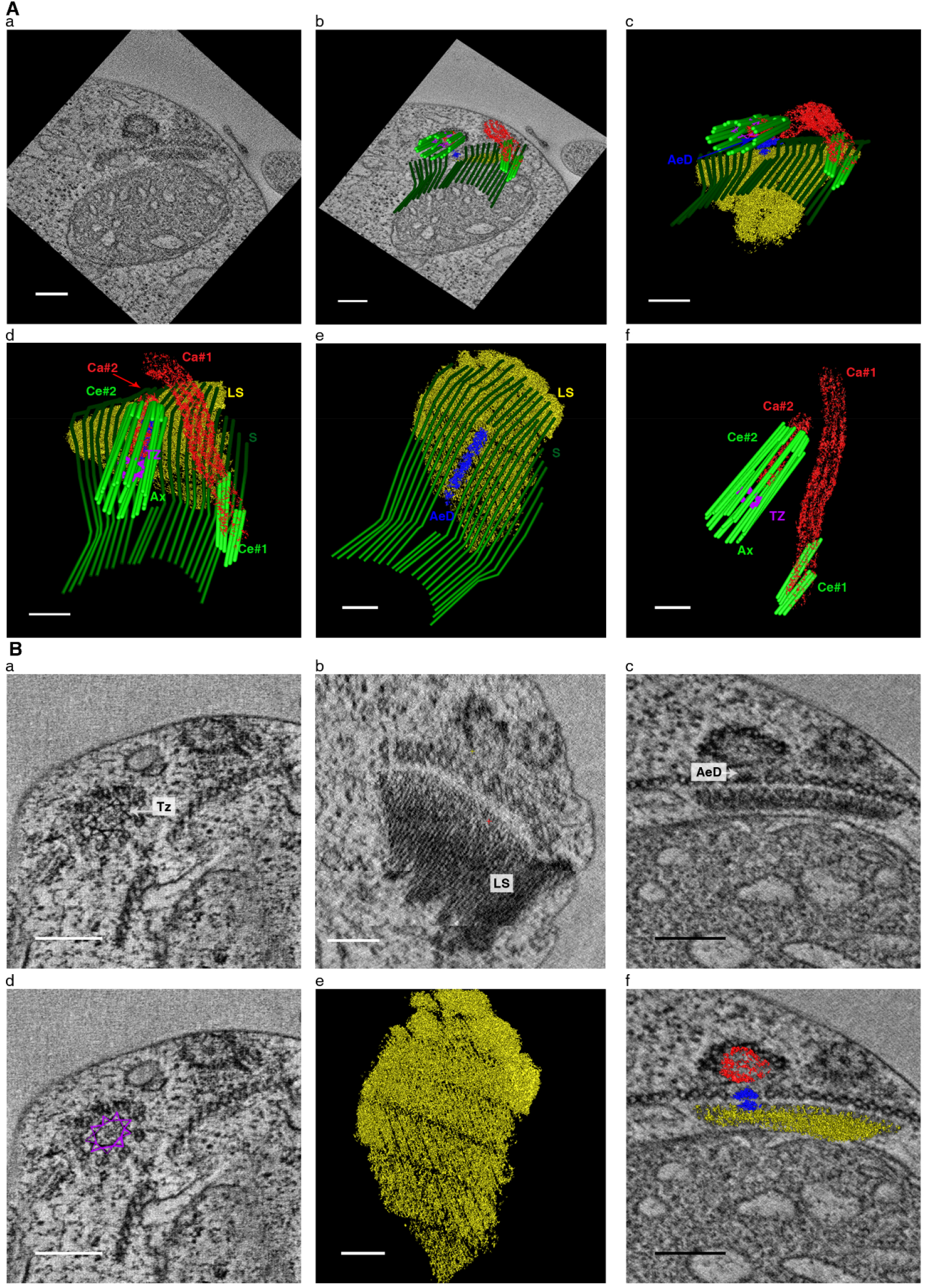
3D analysis of centriole maturation reveals asymmetries between sister centrioles and a long “naked” cartwheel (see movie 3). **A,** From electron tomogram to 3D model. **Aa,** Snapshot of the electron tomogram obtained. **Ab,** Correspondence between the tomogram snapshot and its segmentation. **Ac,** View of the 3D model, highlighting the AeD structure found in one of the centrioles (Ce#2). **Ad,** Top view of the 3D reconstruction is shown in Cc, revealing clear asymmetries between the two sister centrioles. **Ae,** Top view of the MLS, similar to what has been observed in Fig. 2Be. **Af,** The two individual centrioles, revealing clear asymmetries between them. Note that a big portion of one cartwheel (Ca#1) is seen deprived of centriolar microtubules – called here “naked” cartwheel (see also Fig. S1Ba). Furthermore, while one centriolar wall is 9-fold symmetrical (Ce#2), only 2 of the 9 centriolar triplets are observed in the other centriole (Ce#1). Ce #1 and #2 (light green) – wall of centrioles numbered 1 and 2 (arbitrary numbering); Ca #1 and #2 (red) – cartwheel of centrioles number #1 or #2, respectively; LS (yellow) – lamellar strip of the MLS; S (dark green) – spline of the MLS; AeD (blue) – amorphous electron density; TZ (pink) – transition zone (templated from Ce#2); Ax – ciliary axoneme (templated from Ce#2). **B,** Details of the structures identified in the tomogram and model shown in A. **Ba,** Tomogram snapshot highlighting the structure of *P. patens’* transition zone (Tz). **Bb,** Rotated view of the tomogram, highlighting the striation of the lamellar strip (LS). **Bc,** Tomogram view of the AeD connecting one centriole (Ce#2) to the LS. **Bd,** Superimposition of the Tz model with the tomogram snapshot displayed in Ba. **Be,** Top view of the LS reconstruction, highlighting its striated organization. **Bf,** Superimposition of the 3D model with the view shown in H, highlighting the connection between the LS, AeD and one of the cartwheels (Ca#2). Scale-bars = 200nm.

### 3D analysis of centriole maturation reveals intermediate stages and novel structures

In ET, the information collected through passing a beam of electrons through the sample tilted at incremental degrees is used to assemble a three-dimensional image of the target.

#### Centrioles assemble as a bicentriole MTOC and then split into separate centriole units

The bicentriole is an intriguing structure which has only been seen associated with *de novo* biogenesis and for which little detail is known (Moser and Kreitner, 1970; Robbins, 1984). We were able to capture the bicentriole by ET (Fig. 2A, movie 1). Its segmentation (3D model) confirms previous observations that this structure comprises two similar 9-fold symmetrical centrioles (Fig. 2Aa-d) with triplet microtubules (around 200-250nm long), arranged linearly (Fig. 2Af) and connected by a common cartwheel (Fig. 2Ae). Furthermore, we also observed that several individual MTs emanate from this bicentriole, suggesting it functions as an MTOC (Fig. 2Ab-c).

We were able to capture the stage after bicentriole splitting and before ciliogenesis (Fig. 1Cc), where centrioles were oblique to each other and both were anchored to the MLS (Fig. 2B, movie 2). In our 3D model (Fig. 2Bc-d), one centriole (Ce#1) appeared to be slightly longer (approx. 750nm) than the other (approx. 550nm), suggesting that centrioles elongate independently. At this stage both centrioles contained MT triplets of slightly different sizes and a cartwheel structure throughout their entire length (Fig. 2Bf). Below the centrioles and the spline, the striation of the LS was apparent (Fig. 2Ba). Moreover, a gap between adjacent MTs was seen within the spline, under one of the centrioles. This gap was occupied by a previously uncharacterized amorphous electron density (AeD) that localizes between the LS and the centriole (Fig. 2Bd-e).

#### Sister centrioles mature asymmetrically

Centriole maturation to basal body involves the formation of a transition zone, an area at the distal part of centriole which connects the ciliary microtubule structure (the axoneme) with the membrane, while controlling molecule transport into this axoneme.

The structures observed at this stage were very intriguing, showing a remarkable asymmetry and a unique architecture (Fig. 3, movie 3). We observed that both cartwheels outgrew from the centriole walls without any MTs attached – from here onwards called “naked” cartwheels, with one of them extending much further (Fig. 3Ad). This was unexpected given that MTs are thought to provide stability to the cartwheel. The shortest centriole (Ce#2) showed the shortest cartwheel (Ca#2) that was decorated by a full 9-fold symmetrical centriolar wall (Fig. 3Ad, 3Af). A naked cartwheel was found at the proximal end; at the distal end, a transition zone connected the basal body to the membrane and the axoneme (Fig. 3Ba, 3Bd). This transition zone showed a similar structure as observed for *Chlamydomonas reinharditii* (O’Toole *et al*., 2003) and *Marchantia polymorpha* (Carothers and Kreitner, 1968) encompassing the characteristic stellate fiber pattern (Fig. 3Ba, 3Bd).

The centriole with the longest naked cartwheel (Ca#1) showed only two MT triplets throughout the depth of the tomogram obtained (around 3.3μm) (Fig. 3Af), suggesting that some MT triplets might elongate further than others. No major alterations were detected in terms of the organization of the multilayered structure in this stage of spermatogenesis in *P. patens*. The previously described amorphous electron density (AeD) was observed in close contact with the LS and the shortest cartwheel (Fig. 3Bc, 3Bf), while the lamellar strip showed its characteristic striations (Carothers and Duckett, 1980) (Fig. 3Bb, 3Be).

Together, our observations show that centrioles assembled from the same bicentriole (sister centrioles) are initially identical, however upon splitting and maturation, they elongate asymmetrically. This centriole asymmetry indicates that cartwheel and possibly centriole length could be, at least temporarily, distinct between the two sister centrioles. Furthermore, different MT triplets seem to have different lengths along the same mature centriole, in concordance with previous observations in *Marchantia polymorpha* (Carothers and Kreitner, 1968). This surprising asymmetry raises important questions as to how distinct features are generated from seemingly similar entities, in particular, what molecular processes regulate the elongation of the naked cartwheel and/or specific MT triplets, and whether there is a functional consequence of such differences. We next investigated the molecular processes underlying the assembly of the different structures.

### Localization of the evolutionary conserved proteins SAS6 and POC1 shows centriole asymmetry at the molecular level

A set of centriolar components comprising SAS6, BLD10 (also known as CEP135) and POC1, amongst a few others, have been identified as evolutionary conserved across ciliated species, including *P. patens* (Carvalho-Santos *et al*., 2010; Hodges *et al*., 2010). Using a gene expression atlas generated by us and publicly available in the EVOREPRO database (www.evorepro.plant.tools (Julca *et al*.)), we have determined these genes to be preferentially expressed during spermatogenesis, suggesting they could play a role in *de novo* centriole biogenesis (Fig. S2A).

To investigate the assembly of the long cartwheel, as well as the asymmetric centriole wall, we focused on the major core component of the cartwheel – SAS6 (Gopalakrishnan *et al*., 2010; Guichard *et al*., 2017; Nakazawa *et al*., 2007), as well as POC1, which is known to be involved in centriole length regulation (Keller *et al*., 2009). In order to study the localization of these proteins, we generated transgenic reporter plant lines, knocking in tags through homologous recombination guided by CRISPR (see Methods and Table S2 for more details). Given the small size of the structures and difficulty in resolving them by conventional light microscopy, we characterized the localization of SAS6-mcherry and POC1-Citrine during spermatogenesis by 3D-Structured Illumination Microscopy (3D-SIM) imaging of isolated developing sperm cells from these transgenic lines (Fig. 4A).

**Figure 4.**
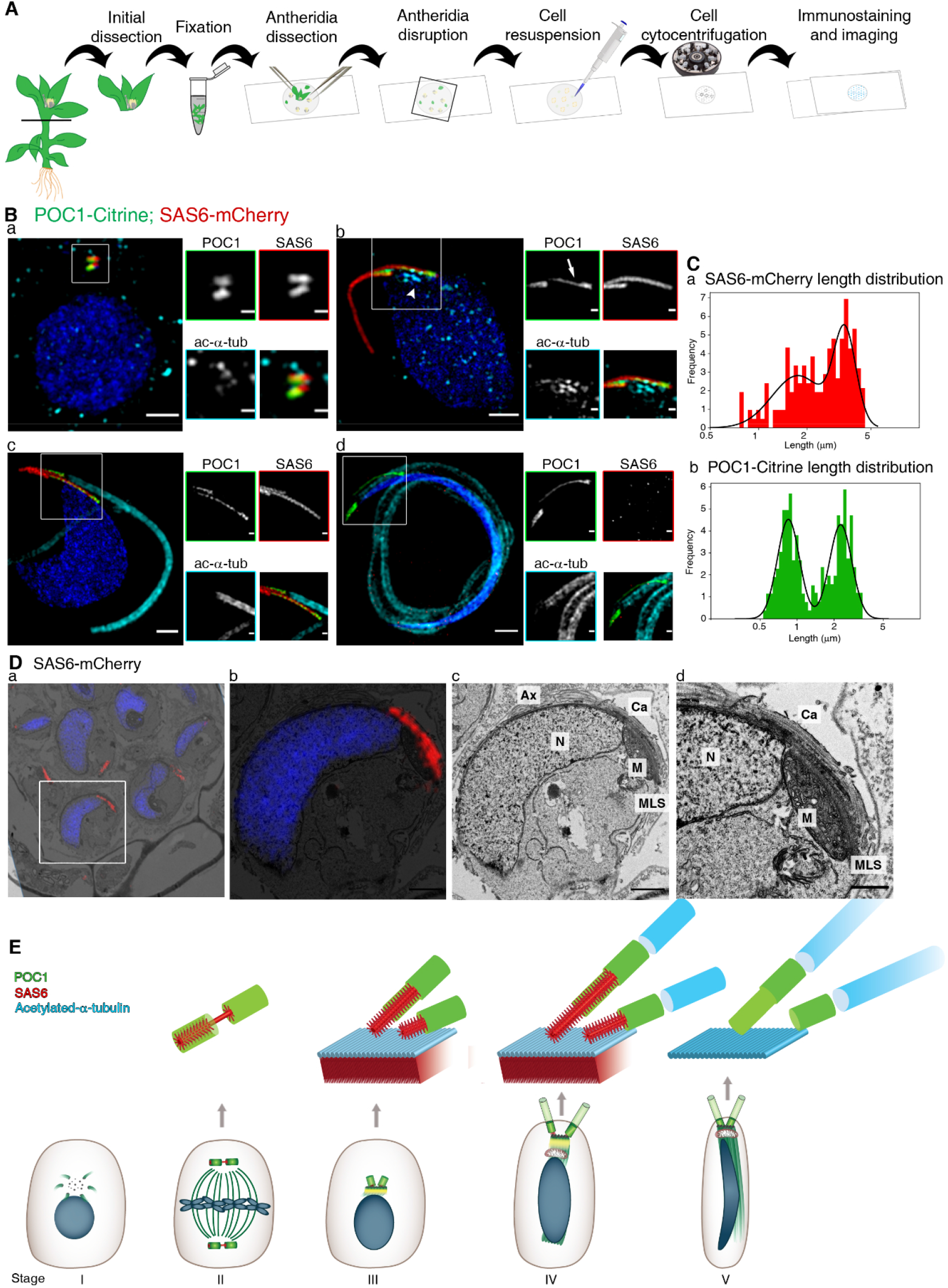
SAS6 and POC1 localization throughout centriole assembly stages supports their role in centriole assembly, elongation and structural asymmetry. **A,** Sample preparation for 3D-SIM and cell stages considered. The top portion of the plants were dissected and fixed. Individualized antheridia were physically disrupted, releasing the developing sperm cells. These were centrifuged onto a coverslip and then stained and imaged (see methods). **B,** Representative 3D-SIM images of individualized cells at different developmental stages from POC1-Citrine; SAS6-mCherry plants. Developmental stages were differentiated based on round nucleus (**Ba**), nuclear elongation without cilia (**Bb**), nuclear elongation with cilia (stained by acetylated-α-tubulin, **Bc**) and condensed and thin nuclear shape (**Bd**). Note the acetylated tubulin signal in the spline (Bb arrowhead) and the elongation of a thinner portion of one POC1 signal (Bb arrow). Blue – DAPI; Green – POC1-Citrine; Red – SAS6-mCherry. Scale-bars = 1μm or 250nm (insets). For sample size please refer to Table S1. **C,** SAS6 and POC1 length distribution (in log scale) reveals the existence of two distinctive centriole populations. **Ca,** SAS6-mCherry length distribution (red) and corresponding Gaussian fitting for both curves (black line, see Fig.S2Ba). Measurements from 75 cells (stages III. and IV.). **Cb,** POC1-Citrine length distribution (green) and respective Gaussian fitting of the two distributions (black line, see Fig.S2Bb). Measurements from 100 cells (stages III. to V.). **D,** Correlative light and electron microscopy (CLEM) of SAS6-mCherry (red signal) sperm cells show its localization at the cartwheel and MLS. **Da,** Alignment of TEM and confocal microscopy images, highlighting the selected cell. Cells were aligned based on nuclear shape (stained with DAPI - blue). **Db,** Overlap of confocal and TEM images from the selected cell in Da. **Dc,** TEM image of the selected cell in Db. **Dd**, Higher magnification TEM image of the SAS6-containing structures from Db. Ax – axoneme; Ca – cartwheel; M -mitochondrion; MLS – multilayered structure; N – nucleus. Scale-bars = 1μm (Dc), 500nm (Dd). **E,** Final scheme of the bicentriole pathway for centriole biogenesis and maturation using the information from EM (Fig. 1–3) and 3D-SIM (Fig. 4B) data. This revealed the existence of long portions of naked cartwheel (red) and the asymmetrical elongation of the centrioles (green), with some microtubules elongation further within the same centriolar wall. Cyan - acetylated-α – tubulin; Green - POC1; Red - SAS6. Note the existence of two bicentrioles at the spindle poles is supported by Fig. S3, while the disappearance of SAS6/cartwheel in stage V is supported by data from 3D-SIM (Fig 4Bd) and EM (Fig. S1B).

In young sperm cells both proteins, SAS6 and POC1, co-localize in two parallel bars near the round nucleus. These structures also appear to contain some acetylated tubulin (Fig. 4Ba), possibly representing the centrioles or bicentrioles. Since we detected two structures, each containing two “dots” of the centriolar components SAS6 and POC1 (Fig. S3A), it is likely that two bicentrioles assemble in the sperm mother cell. Furthermore, these bicentrioles localize in the spindle poles during mitosis (Fig. S3B and 4E), suggesting that their positioning at the poles ensures that each daughter cell inherits only one bicentriole and forms only two cilia.

As spermatogenesis proceeds, SAS6-mCherry and POC1-Citrine signals elongate (Fig. 4Bb-c). A broad and faint tubulin acetylation signal appears along one side of the elongating nucleus (Fig. 4Bb arrowhead), which may represent the spline, as it is made of microtubules. Furthermore, clear tubulin acetylation signal can be also observed on both ciliary axonemes (Fig. 4Bc-d, 4E). However, elongation is asymmetrical between the two originally identical bars, yielding distinct signal lengths of SAS6 and POC1 (Fig. 4Bb-c, 4E). To further understand the consistency of this asymmetry, we have measured the length of SAS6 and POC1 fluorescence signals. Analysis of the signal length distribution has revealed the existence of two distinct populations both for SAS6 (average of 1.8μm and 3.3μm; Fig. 4Ca, S2Ba) and POC1 (average length of 0.8μm and 2.2μm; Fig. 4Cb, S2Bb). Importantly, classification of pairwise lengths per cell into these populations has revealed that 71% (54/76) of cells contain one long and one short SAS6-mCherry signal, while this occurs in 94% (94/100) of cells containing POC1-Citrine. These results strongly support the existence of two centrioles with different lengths in each cell, as well as that there is part of the centriole which is both SAS6 positive and POC1 negative, likely corresponding to the observed naked cartwheels.

It is noteworthy that only a thinner portion of the POC1 bar elongates (Fig. 4Bb POC1 inset arrow), contrary to what we observed for the SAS6 signal (Fig. 4Bb-d). As POC1 in *Chlamydomonas* was suggested to localize to the A-C linker and inner scaffold of the centriole wall (Le Guennec *et al*., 2020; Li *et al*., 2019), this partial elongation of POC1 signal may be associated with the longer microtubule triplets within a centriole, in agreement with our 3D model (Fig. 3). Given that POC1 always localized to the tip of the SAS6 signal from where cilia were nucleated (see acetylated tubulin signal), we propose that the naked cartwheel localizes to the proximal part of the centriole. While POC1 signal remained during final cell maturation (stage V), SAS6 signal disappeared, which correlated with the apparent absence of a cartwheel seen by EM (Fig. 4Bd, 4E, S1B).

Surprisingly, our 3D-SIM imaging of protein localization throughout spermatogenesis revealed three SAS6-mCherry filaments per cell (Fig. 4Bc, 6Ab), while POC1 (Fig. 4Bc) and cilia (Fig. 4Bc, 6Ab) were only seen protruding from two of those. SAS6 is a major cartwheel component, yet we observed only two cartwheels in *P. patens* (Fig. 1–3). This, together with the observation that SAS6-mCherry signal (Fig. 4Bd), cartwheels (Fig. S1B) and the LS (Fig. 1Bf), all disappear upon sperm cell cytodifferentiation, led us to hypothesize that SAS6 could localize to the MLS besides the cartwheels. To test this hypothesis, we performed correlative light and electron microscopy (CLEM), using SAS6-mCherry plants (Fig. 4D). Intriguingly, we observed SAS6 localization to the cartwheels, as expected, but also to the MLS (Fig. 4D, 4E). To our knowledge SAS6 has only been seen associated to the cartwheel in other species. Localization of SAS6 in *P. patens* to both elongated cartwheels and to the plant-specific MLS, suggests putative new roles for SAS6 in land plants.

Collectively, our results support the presence of novel structures, such as naked cartwheels, and structural asymmetries between the two sister centrioles. Hence, centrioles are assembled as similar entities (Fig. 2, 4Ba, 4E) yet mature asymmetrically (Fig. 3, 4Bc-d, 4C, 4E). Given that both SAS6 and POC1 signals (Fig. 4E) localize to the structurally asymmetric regions (Fig. 3) and have known roles in cartwheel assembly (Gopalakrishnan *et al*., 2010; Guichard *et al*., 2017; Nakazawa *et al*., 2007) and centriole elongation (Keller *et al*., 2009) respectively, we speculated whether these proteins could have maintained the same core function while tailoring it to the specificities encountered in *P. patens*. To investigate the functions of those proteins in centriole assembly and maturation, we employed a reverse genetics approach, generating null mutants.

### *P. patens* SAS6, BLD10 and POC1 have conserved functions

#### Despite localizing to both cartwheels and to the multilayered structure, SAS6 is only required for cartwheel assembly

In order to address SAS6’s functions in *P. patens*, we generated SAS6 K.O. (*Δsas6,* Fig. 5B) plants. As *γ*-tubulin is a known component of both the centrosome (Stearns *et al*., 1991) and plant acentrosomal MTOCs (Shimamura *et al*., 2004), we used a *γ*-tubulin2-Citrine (Nakaoka *et al*., 2012) *P. patens* strain as genetic background in our experiments (Fig. 5A) to generate null mutants of the conserved centriole components. Unexpectedly, we found that *γ*-tubulin2 does not localize to the base of the cilia, where centrioles/basal bodies are (Fig. 5Ab). Furthermore, when SAS6-mCherry was introduced into this strain, *γ*-tubulin2 *foci* were found to localize in a non-overlapping manner below the SAS6 signal (Fig. 6Ab). Therefore, we conclude that in *P. patens* sperm cells *γ*-tubulin2 does not co-localize to the centrioles. In agreement with this, none of the knockout (K.O.) lines analyzed in this work displayed significant alterations in *γ*-tubulin2 localization (Fig. 5Bb and 6Bb, 6Cb).

**Figure 5.**
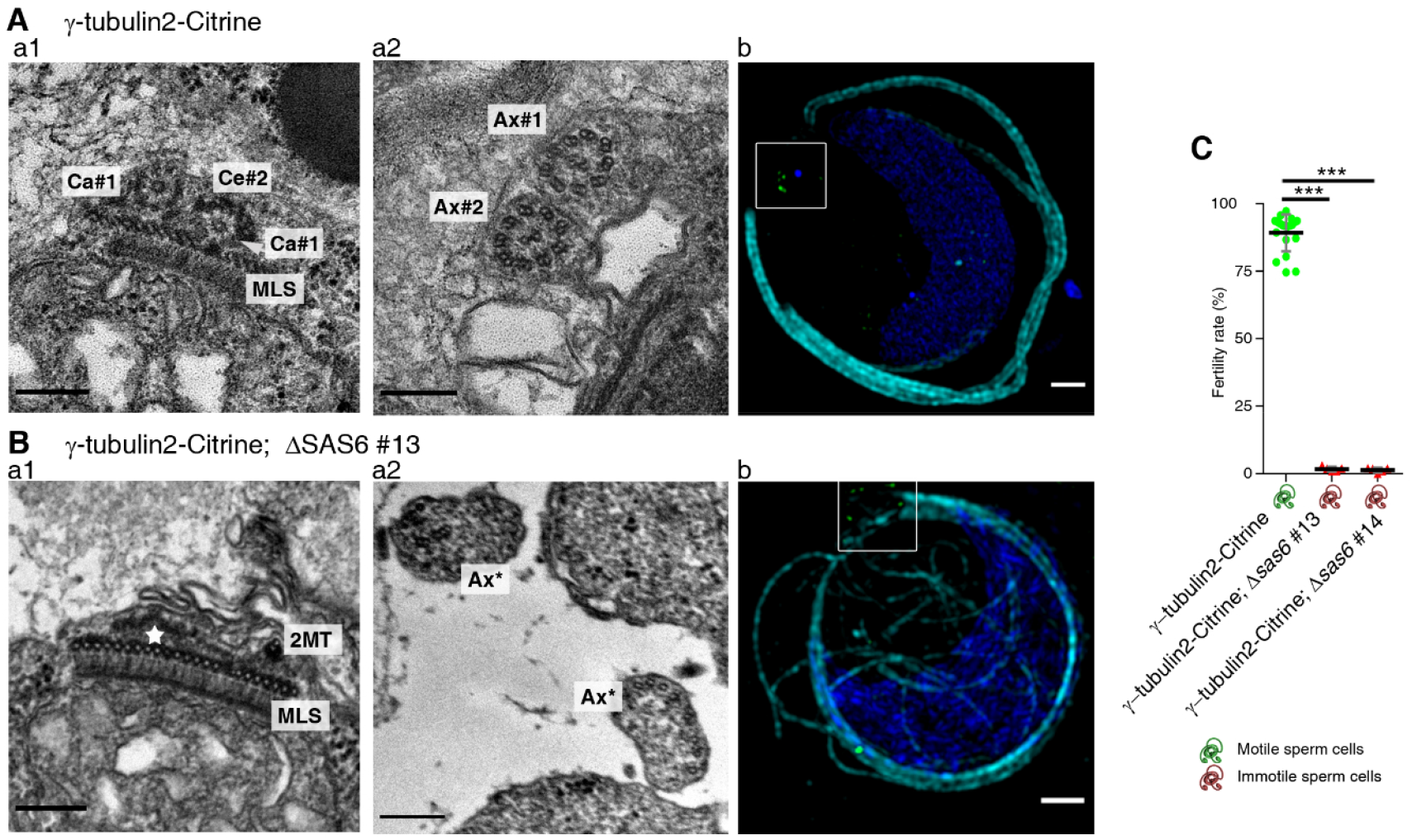
SAS6 is only essential for cartwheel assembly. **A,** γ-tubulin2-Citrine sperm cells (genetic background for *Δsas6).* **Aa,** TEM analysis revealing centrioles docked to MLS (**Aa1**) and the 9+2 axonemal microtubule arrangement (**Aa2**). Ca#1 and #2 - cartwheels of different centrioles (arbitrary numbering); Ce#1 and #2 - centriolar walls around cartwheels #1 and #2, respectively; MLS - multilayered structure; Ax#1 and #2 - axonemal structures #1 and #2 (arbitrary numbering). Scale-bars = 200nm. **Ab**, 3D-SIM of a developing sperm cell, showing the presence of two cilia. Cyan – acetylated-α-tubulin; Green - γ-tubulin2-Citrine; Blue – DAPI. Scale-bars = 1μm or 250nm in insets. **B,** γ-tubulin2-Citrine; *Δsas6* plants. **Ba,** TEM images show the absence of any recognizable cartwheel structure and presence of the MLS (**Ba1**) and disorganized cilia-like structures (**Ba2**). 2MT - microtubule doublet; star - concentrated electron dense material; MLS -multilayered structure; Ax* - disorganized microtubule filaments. Scale-bars = 200nm. **Bb**, 3D-SIM showing aberrant microtubule organization. Cyan – acetylated-α-tubulin; Green - γ-tubulin2- Citrine; Blue – DAPI. Scale-bars = 1μm or 250nm in insets. **C,***Δsas6* plants are sterile. Fertility rate of *Δsas6* (red triangles) compared to its genetic background (γ-tubulin2-Citrine, green dots). Bars represent the mean ± s.d. Significance of differences assessed by one-way ANOVA and Tukey’s multiple comparison test. *** *p-value* < 0.001. The green or red sperm cells symbol indicate motile or immotile sperm cells, respectively (see movie 4). For sample size considerations see Table S1.

**Figure 6.**
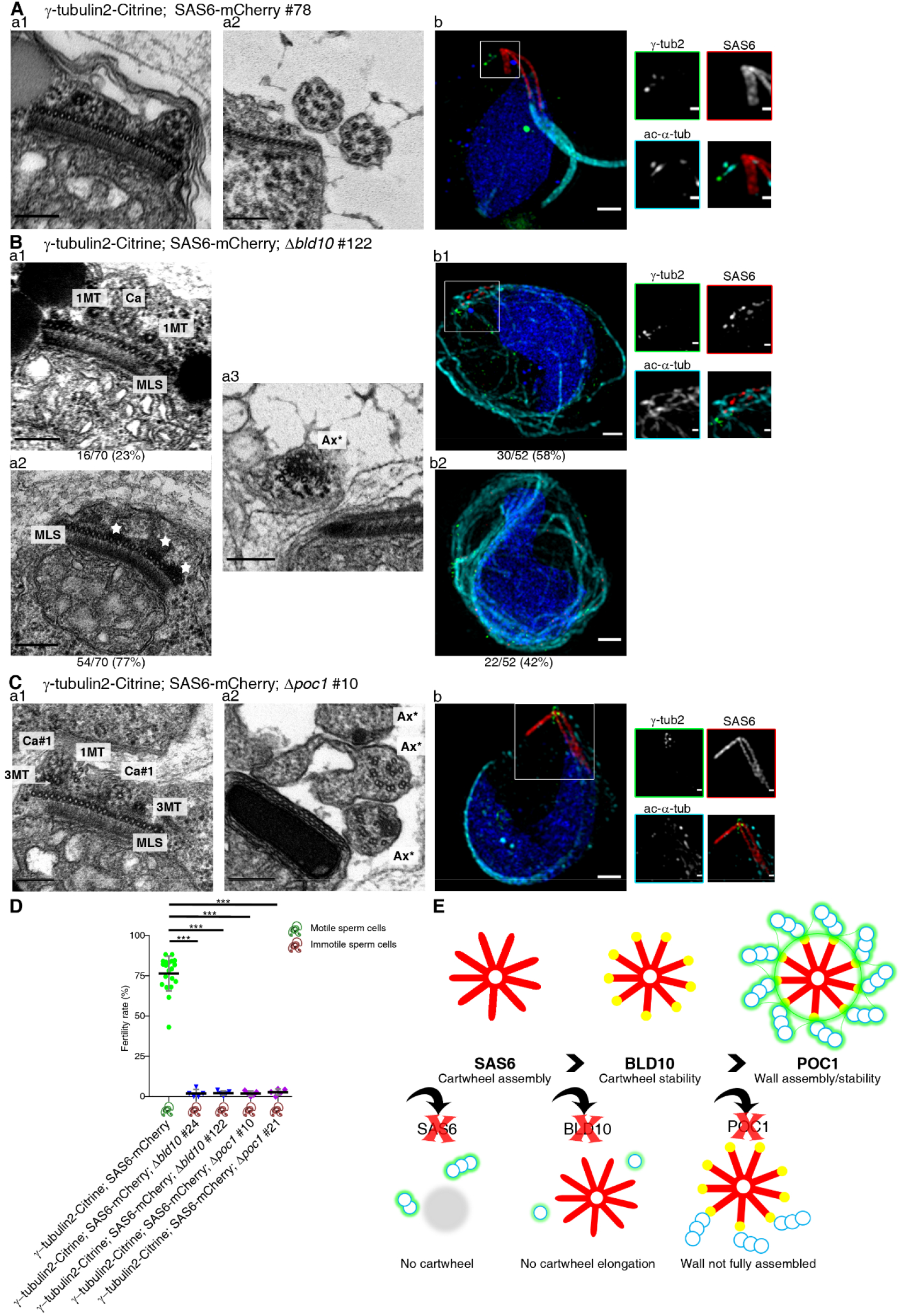
BLD10 and POC1 are important for cartwheel stability and centriole wall elongation. **A,** γ-tubulin2-Citrine; SAS6-mCherry spermatogenesis analysis (genetic background for *Δbld10* and *Δpoc1* plants). **Aa,** TEM captions displaying the sister centrioles docked to the MLS (**Aa1**) and the 9+2 axonemal structure (**Aa2**). Ca#1 and #2 - cartwheels of different centrioles (arbitrary numbering); Ce#1 and #2 microtubular walls around cartwheels #1 and #2, respectively; MLS - multilayered structure; Ax#1 and #2 - axonemal structures of cilia #1 and #2 (arbitrary numbering). Scale-bars = 200nm. **Ab,** 3D-SIM of developing sperm cells showing elongated SAS6 signals and two cilia stained. Cyan – acetylated-α-tubulin; Green - γ-tubulin2-Citrine; Red - SAS6-mCherry; Blue – DAPI. Scale-bars = 1μm or 250nm in insets. **B,** γ-tubulin2-Citrine; SAS6-mCherry; *Δbld10* genotype. **Ba,** TEM reveals that when cartwheel rings are detectable (23% MLS-containing sections), microtubules are not bound to them (**Ba1**). However, more often (77%) no cartwheel rings are observable above the MLS (**Ba2**), with electron dense material concentrated in this region (stars). Moreover, MT-containing protrusions are rarely found and severely disorganized (**Ba3**). 1MT - microtubule singlet; Ca - cartwheel stack; MLS - multilayered structure; Ax* - disorganized microtubule bundles. Scale-bars = 200nm. **Bb,**3D-SIM analysis showing that, while some cells (58%, **Bb1**) have compromised SAS6-mCherry signal (red) elongation, others (42%, **Bb2**) completely lack such signal. In both cases, disorganized microtubule filaments are observed (cyan) and no SAS6 signal is observed in the MLS. Cyan – acetylated-α-tubulin; Green - γ-tubulin2-Citrine; Red - SAS6-mCherry; Blue – DAPI. Scale-bars = 1μm or 250nm in insets. **C,** γ-tubulin2-Citrine; SAS6-mCherry; *Δpoc1* plants. **Ca,** TEM images with observable cartwheel rings with some triplet MT attached (**Ca1**). Accordingly, axonemal symmetry is also incomplete (**Ca2**). 1MT - microtubule singlet; 3MT - microtubule triplet. Ca#1 and #2 cartwheels of different centrioles (arbitrary numbering); MLS - multilayered structure; Ax* - incomplete axoneme. Scale-bars = 200nm. **Cb,** 3D-SIM image displaying a normal SAS6 signal but abnormal microtubule organization. Cyan – acetylated-α-tubulin; Green - γ-tubulin2-Citrine; Red - SAS6-mCherry; Blue – DAPI. Scale-bars = 1μm or 250nm in insets. **D,** Fertility rate of *Δbld10* K.O. (blue inverted triangles) and *Δpoc1* (pink diamonds) plants, as well as their genetic backgrounds (γ-tubulin2-Citrine; SAS6-mCherry, green dots). Bars represent the mean ± s.d. Significance of differences assessed by one-way ANOVA and Tukey’s multiple comparison test. *** *p-value* < 0.001. The green or red sperm cells symbol indicate motile or immotile sperm cells, respectively (see movie 4). For considerations on sample size refer to Table S1. **E,** SAS6, BLD10 and POC1 functions which are conserved across evolution. SAS6 (red) is the major component of the cartwheel, while BLD10 (yellow) is required to ensure cartwheel elongation/stability and microtubule attachment. POC1 (green) is required for complete centriolar wall assembly/stability. Mutant phenotypes highlighted represent our observations in *P. patens*.

We proceeded to investigate *Δsas6* plants using TEM. We first observed that cartwheels were not assembled, although electron-dense material still concentrated above the spline, where the cartwheel was seen in wild-type cells (Fig. 5Ba1 star). Despite SAS6 being localized to the MLS (Fig. 4D), no structural defects were visible in this structure in *Δsas6* (Fig. 5Ba1). Interestingly, despite the lack of cartwheels, abnormal microtubule bundles were assembled in axoneme-like structures, suggesting the presence of cartwheel independent mechanisms to generate organized microtubule bundles (Fig. 5Ba2, 5Bb). Concordantly, individual MT doublets and triplets could be frequently found above the MLS in *Δsas6* sperm cells (Fig. 5Ba1). Moreover, POC1 localization was affected by SAS6 deletion, with several small and discontinuous *foci* being visible in multiple (66%, Fig. S4Ba) or one (34%, Fig. S4Bb) region of the cell. Therefore, *P. patens* SAS6 is essential for the formation of the long cartwheels, however it plays no significant role in the assembly of the plant specific MLS (Fig. 6E).

#### BLD10 and POC1 are involved in cartwheel elongation and centriolar wall assembly, respectively

Given the conserved role of *P. patens* SAS6 in the formation of the long cartwheels, we asked how this structure is stabilized. In several species, BLD10/CEP135 is a known component of the cartwheel spokes required for cartwheel stability (Guichard *et al*., 2017; Hiraki *et al*., 2007; Jerka-Dziadosz *et al*., 2010; Matsuura *et al*., 2004). *Bld10* is also present in the *P. patens* genome and is expressed during spermatogenesis (Fig. S2A). We generated BLD10 K.O. (*Δbld10*) cells in *γ*-tubulin2-Citrine; SAS6-mCherry background. In these mutants, cartwheel rings were only visible in 23% of the analyzed MLS-containing sections. Interestingly no MTs were attached to these cartwheel stacks (Fig. 6Ba1), while singlets or doublet MTs, but never triplets were observed in the cytoplasm of *Δbld10* sperm cells. Despite the absence of any cartwheel-like structures (Fig. 6Ba2), normal MLS with abnormal electron-dense material concentrated above the spline were frequently (77%) observed.

Given the previously observed SAS6 localization at the MLS, we asked whether BLD10 is important to stabilize this SAS6 pool. Indeed, SAS6 signal elongation was compromised in *Δbld10*, with most (58%) sperm cells displaying several small discontinuous SAS6-mCherry *foci* (Fig. 6Bb1) as opposed to the long and continuous SAS6 filaments observed in control cells (Fig. 6Ab). Yet, in the remainder cells analyzed (42%), no SAS6 signal could be detected (Fig. 6Bb2). Interestingly, in a different *Δbld10* line, where POC1 was labelled, we observed POC1 signal in close proximity to the fragmented SAS6 signal (Fig. S4Da), while a dispersed POC1 signal was still detectable in the absence of SAS6 signal (Fig. S4Db). These results suggest BLD10 is important to stabilize SAS6, independently of its localization, while POC1 may be binding to the singlet and doublet microtubules observed in *Δbld10*.

Intriguingly, microtubules that wrap along the cell were observed by 3D-SIM imaging, being more prominent in cells without visible SAS6 signal (Fig. 6Bb2, S4Db). To investigate whether axonemes were present in those cells we used TEM. We observed axonemal-like structures which were severely disorganized and only contained single MTs (Fig. 6Ba3). Therefore, BLD10 appears to have a conserved role in cartwheel/SAS6 stability in *de novo* bicentriole biogenesis (Fig. 6E). Moreover, *P. patens* BLD10 may also be involved in cartwheel-microtubule attachment and in triplet formation, as previously reported in *Chlamydomonas* (Pearson *et al*., 2009) and *Paramecium* (Meehl *et al*., 2016). On the other hand, our results suggest that POC1 has a more downstream role, being directly bound to the microtubules (Fig. 4B, 4D), which is consistent with its reported functions in centriole length and stability in *Chlamydomonas reinhardtii*, *Tetrahymena thermophila* and human cells (Keller *et al*., 2009; Pearson *et al*., 2009; Venoux *et al*., 2012).

Given the distinct size of the two sister centrioles and the seemingly different length of MT triplets around the same centriolar wall, we wondered if *P. patens’* POC1 could be involved in the regulation of centriole MT triplet length. In contrast to the severe *Δsas6 and Δbld10* phenotypes, POC1 K.O. (*Δpoc1,* Fig. 6C) sperm cells assembled normal SAS6-containing cartwheels, capable of elongation (Fig. 6Ca1, 6Cb). However, no more than 3 MT triplets were ever found simultaneously attached to a cartwheel ring (Fig. 6Ca1). Moreover, the MTs attached to the cartwheels were always located close to the spline, suggesting this structure may have a stabilizing effect on the microtubules. Consequently, ciliary axoneme symmetry was incomplete (Fig. 6Ca2). Therefore, our data supports *P. patens* POC1’s critical role in the assembly/stability of the 9-fold symmetrical centriolar wall (Fig. 6E).

In agreement with the ciliary defects observed in all the K.O. genotypes evaluated in this study (*Δsas6, Δbld10* and *Δpoc1*), sperm cell motility (movie 4) and consequently plant fertility (Fig. 5D, Fig. 6D) were severely affected in the absence of precise centriolar structures.

To conclude, the investigated mutant genotypes led to impaired cartwheel formation and stability (*Δsas6* and *Δbld10*) and MT wall assembly and/or stability (*Δpoc1* and *Δbld10*), while leaving structures such as the MLS intact. Overall our data supports the hypothesis that SAS6, BLD10 and POC1 are part of an evolutionary conserved module required for centriole biogenesis (Fig. 6E). Nevertheless, given the observed localization and mutant phenotypes it is also likely that these proteins may have species-specific roles in the asymmetrical elongation of naked cartwheels and specific MT triplets within the same centriolar wall.

### *P. patens* cilia can beat asynchronously

We then asked what could be the physiological significance for the unexpected centriole asymmetry seen between the two *P. patens* sister centrioles. In *Marchantia polymorpha*, sperm cells were suggested to have different beating patterns, with the posterior cilium beating in a 3D-lasso pattern while the anterior one displayed only planar beating (Miyamura *et al*., 2002). We hypothesized that the centriole asymmetry observed by us could render the centrioles differentially resistant to mechanical stresses and/or docked at distinct positions/angles on the cell membrane. Both these scenarios could constrain ciliary motility, making the two cilia functionally distinct and promoting distinct beating patterns.

In order to understand if *P. patens’* cilia could beat differently, we imaged isolated sperm cells at a high frame-rate (200 frames per second - FPS) and manually tracked cilia-tip displacement (relative to the nucleus). Our data shows that in 83% (74/89) of the cells we imaged, only one cilium appeared to be actively moving (Fig. 7A, S5A, movie 5; green line), while the inactive one appeared to passively follow the cell’s rotation (Fig. 7A, S5A, movie 5; blue line). By averaging cilia beating patterns from 5 cells it was possible to distinguish the beating cilia, with frequent changes in nuclear distance (Fig. 7Ab green line) from those that appeared to be inactive (Fig. 7Ab blue line). In the remainder 17% of the analyzed sperm cells (15/89) both cilia appeared to be actively moving (Fig. 7B, S5B, movie 6), and no clear differences were detected between the beating pattern of both cilia. These observations indicate that while both cilia are able to beat, most cells show asynchronous beating, with only one cilium appearing to be active.

**Figure 7.**
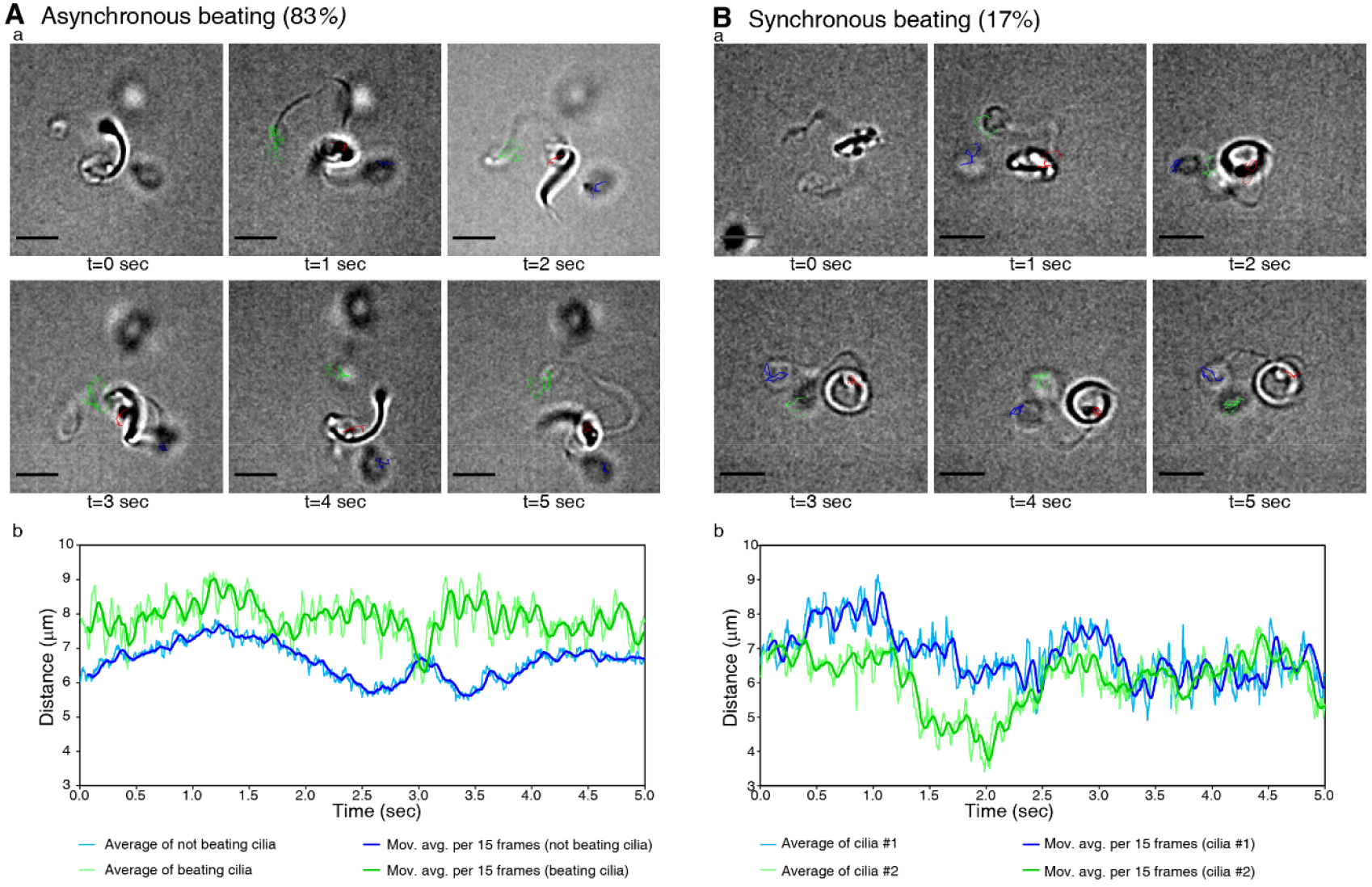
Majority of cells show asynchronous ciliary beating. **A,** Asynchronous ciliary beating. In most sperm cells (83%, 74/89), only one cilium appears to be actively beating (green) while the other (blue) moves only passively (following nuclear movement, red). **Aa,** Snapshots of a sperm cell tracked over time. Scale-bars = 5μm. **Ab,** Ciliary-tip distance across time, showing a distinct pattern between both cilia. Thin lines represent average displacement for the beating cilium (green) or inactive cilium (blue) while thicker and lines represent their respective running averages across 15 frames. Pooled data from 5 cells, for an example of individual cell data see movie 5 and Fig. S5A. **B,** Sperm cells synchronously beating both cilia. Both cilia appear to be active and show similar beating patterns (17%, 15/89). **Ba,** Example of a sperm cell tracked. Scale-bars = 5μm. **Bb,** Ciliary-tip distance over 5 seconds, showing a similar pattern for both (green and blue) cilia. Thin lines represent average displacement for one cilium (green) or the other (blue) while thicker lines represent their respective running averages over 15 frames. Cilia colors were attributed randomly. Pooled data from 5 cells, for an example of individual cell data see movie 6 and Fig. S5B.

## Discussion

Centrioles are widespread in eukaryotes and can be assembled through various structural pathways. Centriole duplication has been extensively studied, however the diverse mechanisms underlying *de novo* centriole assembly have been rarely investigated. Here we took advantage of the tools that exist for the moss *P. patens* to investigate *de novo* centriole biogenesis. Our TEM and ET work delineates a morphological pathway in which a bicentriole structure, composed by two centrioles united by a common cartwheel, might arise in a MTOC, here called concentrator. After bicentriole splitting, the two centrioles mature asymmetrically, through elongation of the naked cartwheel and of specific MT triplets (Fig. 1–3). These asymmetric centrioles template the assembly of two motile cilia, which often beat asynchronously (Fig. 7). Remarkably, despite the presence of different structures in *P. patens*, we show that evolutionary conserved proteins present in all branches of the eukaryotic tree of life, also have essential structural roles in bicentriole-mediated *de novo* centriole assembly (Fig. 4–6).

### Temporal, spatial and numerical regulation of centriole assembly

Centriole duplication is known to be tightly regulated in number (one daughter per mother centriole), time (cycle-cycle coupling), and space (daughters assemble orthogonally to their mothers) (Nigg and Holland, 2018). Polo-like kinase 4 (PLK4) is a master regulator of centriole duplication as its concentration and activity control centriole biogenesis in most animal cells (Bettencourt-Dias *et al*., 2005; Habedanck *et al*., 2005; Kleylein-Sohn *et al*., 2007). However, PLK4 orthologs have not been found outside of opistokhonts (Carvalho-Santos *et al*., 2010; Hodges *et al*., 2010). Therefore, other molecules and mechanisms may regulate centriole biogenesis. We propose that the timing of centriole biogenesis in *P. patens* might be defined by the specific transcription of conserved centriolar components, as they appear to be preferentially expressed during spermatogenesis (Fig. S2A).

What could regulate the localization and number control of *de novo* centriole biogenesis? While during duplication, each mother centriole serves as a platform for daughter centriole assembly, it is unclear how *de novo* biogenesis is spatially regulated. Recent studies highlight the existence and involvement for pericentriolar material in *de novo* assembly in animals (Mercey *et al*., 2019; Nabais *et al*.). Given that many PCM components do not appear to have direct homologs outside of the Holozoa (Hodges *et al*., 2010), it is possible that other molecules with similar function might exist in *P. patens*. In our work, we observed electron dense clouds close to the nucleus, which seemed to nucleate microtubules and to precede centriole assembly (Fig. 1Ba). Furthermore, microtubules also emanated from already formed bicentrioles (Fig. 2A, movie 1). Electron dense clouds which nucleate MTs were also seen at the beginning of *de novo* centriole assembly in the bryophyte *Riella americana* (Robbins, 1984). It is tempting to speculate that microtubule trafficking towards an initially amorphous MTOC might locally concentrate centriolar precursors beyond a critical threshold required for *de novo* centriole assembly.

Finally, a variable number of centrioles is assembled *de novo*, depending on the organism and pathway involved (Nabais *et al*., 2018). In *P. patens*, two bicentrioles assemble in the sperm mother cell (Fig. S3A) and this is also the case in *Riella americana* (Robbins, 1984) and *Marchantia polymorpha* (Moser and Kreitner, 1970), suggesting some general mechanism for number control. The localization of each bicentriole to one pole of the spindle upon mitosis (Fig. S3B), is likely to ensure that each daughter cell only has two centrioles, hence forms two cilia It is possible that the linear cartwheel geometry ensures that only one centriole can assemble at each cartwheel end of the bicentriole, representing a structural mechanism to regulate centriole copy number. In the future, it will be important to understand how cells limit the number of bicentrioles formed.

### Centriole length and structural asymmetry

Centriole structure and size are highly regulated within each cell type in a species. However, much diversity is observed within species and in between species, in terms of centriole symmetry, size and specific features (reviewed in Gupta and Kitagawa, 2018). Despite reaching similar lengths upon maturation, human centriolar microtubule triplets are known to elongate individually during procentriole assembly. The A-tubule grows from its distal-end, while B and C-tubules elongate bidirectionally (Guichard *et al*., 2010). In contrast, cartwheels show proximal-end directional elongation (Aydogan *et al*., 2018) with a small (10-40 nm) portion overhanging from the centriole wall (Klena *et al*., 2020). Indeed, cartwheel and centriolar microtubules appear to assemble interdependently in order to define and stabilize centriole architecture (Hilbert *et al*., 2016).

In this work we reveal that the two sister centrioles assembled from the same bicentriole in *P. patens* are initially identical (Fig. 2A). Yet unexpectedly, they mature asymmetrically. This asymmetry is seen both in terms of cartwheel and microtubule triplet length (Fig. 3, 4), generating two distinct centriole types in the same sperm cell (Fig. 4C). Furthermore, *P. patens* centrioles display elongated naked cartwheels overhanging from their proximal end (Fig. 3, 4B), in agreement with an autonomous proximal-end elongation mechanism (Aydogan *et al*., 2018). Our data also shows that microtubule triplets from the same mature centriole may have distinct lengths (Fig. 3, 4B), a characteristic that maybe more widespread, as also suggested to be present in the liverwort *Marchantia polymorpha* (Carothers and Kreitner, 1968). However, the directionality of such growth remains unclear, possibly requiring live-imaging and/or perturbation experiments. Overall, these features make *P. patens* centrioles an exciting system to study cartwheel and microtubule triplet elongation, and in particular to investigate whether distinct regulatory mechanisms operate in each centriole.

### An evolutionary conserved module in centriole assembly and diversification

Several key centriolar components are known to be conserved in ciliated species (Carvalho-Santos *et al*., 2010; Hodges *et al*., 2010). By exploring their localization (Fig. 4) and function (Fig. 5–6), we show that their critical structural roles in centriole assembly have been mostly conserved throughout evolution. SAS6 and BLD10 are essential for cartwheel assembly and stability respectively, while BLD10 and POC1 appear to be required for assembling the 9-fold symmetrical MT wall. Nevertheless, some specificities exist in the regulation of those components that may underlie novelties. For example, *P. patens’* SAS6 localizes to the MLS and disappears upon cell cytodifferentiation. While cartwheels are known to disappear in human cells during mitosis (Strnad *et al*., 2007), they are retained in the basal bodies of other species (reviewed in Gupta and Kitagawa, 2018).

What elements could lead to diversification of the pathways? It is possible that specific structures, such as the spline and the LS, might play a role in generating asymmetries. The spline was reported to be involved in nuclear elongation in the algae *Nitella* (Turner, 1970). Our work suggests the spline may also stabilize MTs as in *Δpoc1* cells only the MT triplets close to the spline appear to be able to bind the cartwheel spokes (Fig. 6Ca1), being also the longer ones in the asymmetrical centriole walls. The LS contains centrin (Vaughn *et al*., 1993) and SAS6 (Fig. 4D), disappearing upon cell cytodifferentiation (Carothers and Duckett, 1980 and Fig. 1Bf). *γ*-tubulin2 is detected on the opposite side from where cilia grow, most likely the tip of the MLS (Fig. 6Ab). Consequently, we hypothesize that the LS may perform PCM-like functions in concentrating components and/or stabilizing the structures assembled (Pimenta-Marques and Bettencourt-Dias, 2020), such as the naked cartwheel and particular MT triplets. In the future it will be important to be able to manipulate the MLS, as well as to identify PCM-functional equivalents, to test whether these are critical variables in generating diverse structures, such as the ones observed here in *P. patens*.

### Coordination of ciliary beating

*P. patens* motile cilia are required for the sperm cells to swim and reach the egg cell. Based on the centriole asymmetries unraveled in this work (Fig. 3–4) and supported by observations from ciliary beating of the liverwort *Marchantia polymorpha* sperm cells (Miyamura *et al*., 2002), we hypothesized that the two cilia could have distinct beating patterns. We observed that the most common beating pattern was asynchronous, with only one cilium actively beating (Fig. 7A). However, both cilia were able to also beat synchronously, which might lead to the helical motion described by Ortiz-Ramírez *et al*. (2017) (Fig. 7B). *Chlamydomonas* cells grown in the dark can alternate their swimming behavior between periods of synchronous and asynchronous beating, leading respectively to cells moving in a nearly straight trajectory or its reorientation (Polin *et al*., 2009). We envision that external cues (such as signals from the egg) may lead to the alteration in ciliary beating patterns in *P. patens*. In the future it will be important to understand whether asymmetries in basal body structures may influence the regulation of flagella beating.

In this work, we have described with unprecedented detail the bicentriole-mediated *de novo* centriole biogenesis pathway, and how the initially identical centrioles mature into asymmetrical entities, that assemble asynchronously beating cilia. While, we focus on a single species, other land plants have been described to show similar structures, and many organisms form centrioles *de novo*. Our work pioneers the molecular understanding of *de novo* centriole biogenesis and maturation outside of animals, supporting a scenario where centriole biogenesis is less constrained than previously thought. Moreover, our results suggest that the same molecular components can generate diverse structures, even if associated with essential functions such as motility and fertility. Our work highlights the importance of investigating fundamental processes in diverse species as more tools become available.

## Methods

### *P. patens* strains and growth conditions

The Gransden wild-type (WT) *Physcomitrium* (*Physcomitrella*) *patens* strain (Ashton and Cove, 1977) was used in this work. All other plant genotypes were derived from this WT strain or y-tubulin2-Citrine (Nakaoka *et al*., 2012). Detailed information on the genotypes used is available in Table S2.

Plant material was routinely maintained by vegetative propagation of protonema (sub-cultured every 7 days after mechanical disruption (TissueRuptor; Qiagen) on Petri dishes containing KNOPS media55 supplemented with 0.5g/L ammonium tartrate dibasic (Sigma-Aldrich) and grown at 25°C, 50% humidity and 16h light (light intensity 80μlum/m/s) (Fig. 1A). Most experiments required gametangia development. For that, plants were grown for 6-8 weeks at 25°C, 50% humidity and 16h light (light intensity 80μlum/m/s) in Phytatray II (Sigma-Aldrich) containing 4 sterile peat pellets (Jiffy-7, Jiffy Products International). Water was supplied to the bottom of each box. Sexual reproduction (leading to gametangia and sporophyte development) was induced by transferring the Phytatray boxes to 17°C, 50% humidity and 8h light (light intensity 50μlum/m/s) conditions (Fig. 1A). Unless otherwise stated, experiments were performed 15 days after induction (DAI) of sexual reproduction.

### *P. patens* transgenic strain generation

#### Vector construction and preparation

Gene sequences for the conserved centriolar proteins (Carvalho-Santos *et al*., 2010; Hodges *et al*., 2010) SAS6 (Pp3c23_8210v3.1), BLD10 (Pp3c9_9030v3.1) and POC1’s longest CDS (Pp3c16_11590V3.1) were obtained from Phytozome (https://phytozome.jgi.doe.gov/pz/portal.html). Genetic engineering was employed based on homologous recombination (HR) with CRISPR/Cas9 technology being used in specific cases (see Table S2) to create targeted double-strand breaks. In both knockin (fluorescent protein tagging) and knockout strategies, one of the homology arms was the 600bp-1kb sequence downstream of the stop codon. In the knock-in vectors, the other homology arm was the 600bp-1kb sequence before the stop codon, which was immediately followed by a linker and mCherry or Citrine sequences. The other homology arm in the knock-out constructs was the 600bp-1kb upstream sequence to the CDS start codon. In both cases, an antibiotic resistance cassette was introduced between the two homology arms (see Table S1). After molecular cloning of the respective vectors, these were confirmed by endonuclease restriction digestion and Sanger sequencing (GATC) of the engineered regions. The correct HR constructs were linearized, ethanol precipitated and eluted in molecular grade water. The gRNAs for the CRISPR/Cas9 strategy were synthesized as single-stranded oligos (Integrated DNA Technologies), annealed to become double-stranded and then inserted in the same plasmid containing the Cas9 sequence, after sequence confirmation by Sanger sequencing (GATC), these vectors were ethanol precipitated and eluted in molecular grade water. For further details on plasmid engineering, please refer to Table S2.

#### Transformation

5-day old protonema tissue was digested in a 2% Driselase solution (Sigma-Aldrich) in 0.5M D-Mannitol for 2h at RT. Protoplasts were collected by 2 rounds of centrifugation (5min at 250*g*) and filtration (EASYstrainer 40μm, Greiner Bio-One International), followed by a last centrifugation and re-suspension into 3M media (15mM MgCl2; 0.1% MES (w/v) and 0.5M D-Mannitol). 2 million *P. patens* protoplasts (counted in a hemocytometer) were transformed with a total of 30μg of DNA (when several plasmids were used, their amount was equally distributed to a total of 30μg) by careful addition of 300μl 40% PEG-4000 (polyethylene glycol, Serva) in 3M media, followed by incubation at 45°C for 5min. After the heat shock, the transformants were incubated at RT for 15min before careful addition of 10mL of 3M media followed by homogenization. Finally, the transformants were centrifuged (5min at 250*g*) and re-suspended into 3mL of REG media (KNOPS media, 5% Glucose and 3% D-Mannitol). The protoplasts were then transferred to 3.5cm petri dishes (Sarstedt), incubated in the dark overnight (25°C) and regenerated for 10 days at normal growth conditions (25°C, 50% humidity, and 16h light (light intensity 80μlum/m/s).

#### Selection and genotyping

Selection of transformants was achieved by 2-3 rounds of 10-day growth in selective followed by 10-days in non-selective media. Selective media was KNOPS media supplemented with the respective antibiotics: 25μg/mL G418 (Sigma-Aldrich), 30μg/mL Hygromycin B (Alfa Aesar) or 100μg/mL Zeocin (Invitrogen). DNA from surviving plant colonies was extracted by phenol-chlorophorm and PCRs were performed to test for proper genetic integration. Experiments were performed using two independent lines from the same genotype. For further details, see Table S1.

### Transmission Electron Microscopy (TEM)

#### Chemical fixed samples

Top part of individual plants were fixed for 2h at RT in a 6% (v/v) glutaraldehyde (Polysciences) and 0.5% (v/v) Tween 20 in 0.1M phosphate buffer (PB) solution. Samples were then washed twice with 0.1M PB and individually embedded in 2% (w/v) low melting point agarose. The agarose blocks were post-fixed in 1% (v/v) osmium tetroxide in 0.1M PB for 2h on ice. Samples were then washed twice with 0.1M PB and twice with distilled water, followed by en bloc staining with a 1% (w/v) uranyl acetate aqueous solution for 1h at RT. Samples were dehydrated through an ethanol series (30%, 50% for 10 min each; 70% overnight at 4°C; 90% for 10 min at 4°C and 3 times 100% for 15min at 4°C) and infiltrated with increasing concentrations of EMbed-812 epoxy resin (EMS) of 25%, 50%, 75% in ethanol for 90min each). Finally, samples were embedded in 100% epoxy resin which was polymerized at 60°C for 24 h. Ultrathin (70 nm) sections were obtained using a Leica UC7 Ultramicrotome and collected on palladium-cooper grids coated with 1% (w/v) formvar (Agar Scientific) in chloroform. Sections were post-stained with 1% (w/v) uranyl acetate and Reynolds’ lead citrate for 5min each and imaged on a Hitachi H-7650 (100 keV) transmission electron microscope using a XR41M mid mount AMT digital camera. For sample size considerations, please see Table S1.

#### High-pressure frozen-freeze substituted samples, electron tomography and segmentation

Plant top portions were placed in 0.2μm aluminum specimen carriers with 10% (w/v) BSA, 8% (v/v) methanol cryoprotectant solution and frozen using a High Pressure Freezer Compact 02 (Wohlwend Engineering Switzerland). Samples were freeze substituted in a Leica EM AFS2 with a solution of 2% (v/v) osmium tetroxide, 0.2% (w/v) uranyl acetate in methanol, 1% (v/v) distilled water in acetone for 1h at −90°C, followed by a warm-up with a slope of 5°C/h until −80°C. At −80°C samples were incubated for 72h and then warm-up until 0°C where three washes with acetone were done for 10min each. Samples were infiltrated in increasing concentrations (5%, 10%, 25%, 50%, 75% and 100%) of EMbed-812 epoxy resin at RT for at least 4h each. Resin was polymerized at 60°C for 24h. Ultrathin sections of 70 and 300nm were obtained for ultrastructural analysis and tomography acquisitions, respectively. 70nm sections were collected and post-stained as described for chemical fixed samples and imaged with a FEI Tecnai G2 Spirit BioTWIN (120 keV) transmission electron microscope using an Olympus-SIS Veleta CCD Camera. 300 nm sections were collected as described for chemical fixed samples, however and were not post-stained. In these sections, dual-axis tilt series of serial sections were rapidly acquired (Schorb *et al*., 2019) using a Gatan OneView Camera and SerialEM Software (Mastronarde, 2005) on a Tecnai F30 (FEI) operated at 300kV. Electron tomograms were reconstructed from the dual-axis tilt series and serial section tomograms were all stitched in *z* using eTomo/IMOD software (Kremer *et al*., 1996). Some electron tomograms were automatically reconstruction according to (Mastronarde and Held, 2017), while others were individually processed.

#### Correlative light and electron microscopy (CLEM)

Pre-embedding CLEM was performed. Briefly, the top part of SAS6-mCherry individual plants were fixed for 2h at RT using a solution of 2% (v/v) formaldehyde (EMS), 0.2% (v/v) glutaraldehyde, 0.5% (v/v) Tween-20 in 0.1% M phosphate buffer (PB). Then samples were washed three times with PB and incubated with 0.15% (w/v) glycine in distilled water for 10min at RT. Samples were washed three times with distilled water and infiltrated in 30% (w/v) sucrose for cryo-protection ON at 4°C on a rotator. The embedding was made with optimal cutting temperature (OCT) compound (Sakura) before freezing in liquid nitrogen and section using the cryostat (Leica CM 3050 S) slices of 5μm (object temperature −18°C and chamber temperature −22°C). Sections were picked-up in coverslips coated with 2% (v/v) 3-aminopropyltriethoxysilane (Sigma-Aldrich) in acetone and stained with DAPI (Sigma-Aldrich) 1:1000 in PBS for 5min at RT. After washing the sections with PBS three times, the coverslips were mounted with PBS and imaged. The imaging was done using the objectives 20x 0.70NA dry and 63x 1.4NA oil in a confocal microscope (Leica SP5 Live) with DAPI and mCherry signals were excited using 405nm and 561nm lasers respectively. Sections that had signal and were structurally preserved were dismounted from the slide, washed 10 times with PB and post-fixed using a solution of 1% (w/v) potassium ferrocyanide, 1% (v/v) osmium tetroxide in 0.1M PB for 30min at 4°C. Sections were washed twice in 0.1M PB and twice in distilled water before dehydration in an ethanol series of 30%, 50%, 70%, 90% and 100% for 10min each and embedding in 100% EMbed-812 epoxy resin, which was polymerized at 60°C for 24h. Sections were picked-up and post-stain as described for the chemical fixed samples. The low magnification TEM and confocal images were aligned considering DAPI/nucleus shape. Cells with clear elongated SAS6-mCherry signals were selected and serial TEM images were obtained. The confocal images were then superimposed to the serial TEM pictures and the best alignment between DAPI signal and nuclear shape was selected.

### Immunofluorescence and 3D-SIM imaging

#### 3D-SIM

Apical shoots of 10-15 plants were fixed overnight at 4°C with 4% formaldehyde (Polysciences), 1mM MgCl2, 50mM EGTA, 0.2% NP-40 and 1% Triton-X in 1x PBS. Fixed samples were washed twice with 1x PBS. Next, gametangia were carefully dissected under a stereoscope onto a microscope slide and then crushed between the slide and a coverslip. Crushed samples were re-suspended into 100μl of 1x PBS by washing the coverslip and slide with this solution. The solution containing the individualized SCs was then centrifuged (Cytopro cytocentrifuge) for 7min at 500*g* onto a coverslip (Fig. 4A).

The samples’ coverslips were then immunostained. First, nonspecific antibody binding was blocked by incubating the samples at RT for 1h with a blocking solution (5% BSA, 0.1% NP-40 and 0.5% Triton-X in 1x PBS). Immediately afterwards, samples were incubated for 2h at RT with the primary anti-acetylated tubulin antibody (1:1000, Sigma-Aldrich #T7451) in blocking solution, washed three times (10min each) with the blocking solution and incubated for 1h at RT with secondary anti-mouse-cy5 (1:500, Life Technologies A21236) in blocking solution. Next, coverslips were washed with 1x PBS, 0.1% NP-40 and 0.5% Triton-X for 10min. DAPI (1:500, Sigma-Aldrich) was added to this washing solution for a second 20min wash and a last 10min wash (without DAPI) followed. Finally, samples were washed with distilled water for 5min and mounted on slides using vectashield (Vector Laboratories).

3D-SIM imaging was performed on an GE HealthCare Deltavision OMX system, equipped with 2 PCO Edge 5.5 sCMOS cameras, using a 60x 1.42NA Oil immersion objective. Images were reconstructed with Applied Precision’s softWorx software considering a Weiss filter of 0.05. For experimental sample sizes please refer to Table S1.

#### Signals length quantification and distribution

In order to measure SAS6 and POC1 signals, 3D-SIM images from SAS6-mCherry (75 stages III and IV cells) and SAS6-mCherry; POC1-Citrine (100 cells from stages III to V) sperm cells were selected. In these images, the individual channels were split and an auto-Huang threshold applied. Afterwards, automatic ROIs were defined and manually curated to remove regions with overlapping signals from both cartwheels and/or MLS signals. Finally, the longest axis from those ROIs was considered as its length.

In order to test if two distinct length populations of centrioles exist, we fitted a Gaussian mixture model with an increasing number of Gaussian distributions to the data, disregarding the origin of each value. The model with the lowest Bayesian Information Criterion (BIC) was selected as the most parsimonious number of Gaussians (Fig. S2B). This model was then used to predict the class (long or short) of each pair of centrioles in each cell. The number of cells that were classified as having one “long” and one “short” centriole was recorded. Analyses were performed using python’s sklearn package version 0.21.3.

### Sperm cell motility

#### K.O. SC motility assessment

Gametangia from 10-12 individual plants were dissected onto a coverslip containing 50μl of SC media (0.45mM CaCl2, 0.3mM MgSO4, 0.02mM KNO3 and 0.081mM NaHCO3) (Ortiz-Ramírez *et al*., 2017) supplemented with 1μg/mL of the viability dye fluoresceín diacetate (Sigma-Aldrich F7378). Released and viable (checked with the GFP filterset) sperm cells’ clusters were imaged on a commercial Nikon High Content Screening microscope equipped with an Andor Zyla 4.2 sCMOS camera, using the 20x 0.75NA dry objective in the BF channel (33FPS) during 1min. Acquired movie sequences were analyzed using Fiji57 to assess if sperm cells were motile or not (movie 4). Clusters from at least 3 independent experiments were analyzed per genotype (see Table S1).

#### Fast time-lapse acquisition, tracking and data analysis

Gametangia from 5 individual WT plants were dissected onto a coverslip containing 30μl of SC media (0.45mM CaCl2, 0.3mM MgSO4, 0.02mM KNO3 and 0.081mM NaHCO3) (Ortiz-Ramírez *et al*., 2017). Individualized sperm cells from released clusters were imaged on a commercial Nikon High Content Screening microscope equipped with an Andor Zyla 4.2 sCMOS camera, using a 63x 1.27NA water objective in the BF channel (200FPS) during 1min. Acquired image sequences were analyzed using Fiji57. A total of 135 cells were imaged with 89 being classified into two categories based on observed cilia beating patterns (synchronous vs. asynchronous cilia beating). Both cilia’s tip and nucleus from 5 sperm cells from each category were manually tracked (using TrackMate (Tinevez *et al*., 2016) FIJI plugin) for 1000 frames (5 sec). Cilia tip distance was calculated by frame using Euclidean distance in relation to the nucleus.

### Fertility rate analysis and statistics

Assessment of fertility rate was performed as described in ref. (Ortiz-Ramírez *et al*., 2017). Fertility rate is the percentage of gametophores containing sporophytes (Fig 1A), considering at least 100 randomly collected gametophores per sample, 6 weeks after induction of sexual reproduction. Statistical differences were evaluated by one-way ANOVA followed by Tukey’s post-hoc.

## Acknowledgements

We thank Elena Kozgunova (University of Freiburg, Germany) for plants and reagents. We would like to acknowledge the members of the MBD and JDB labs for helpful discussions on the manuscript. We thank Swadhin C. Jana and Paulo Duarte for technical support. We thank the Electron Microscopy Core Facility (EMBL, Heidelberg) for supporting ET data acquisition and Pedro Machado (King’s College London, UK) for support with tomogram segmentation. The authors would like to acknowledge the following IGC facilities: Electron Microscopy (EM protocol implementation and sample preparation), Light Microscopy (for equipment availability), Histopathology (for equipment availability), Quantitative and Digital Science (data analysis and helpful discussions) and Plant facility (plant chamber maintenance).

This work was supported by FCT (Fundação para a Ciência e Tecnologia) fellowship PD/BD/114350/2016 attributed to S.G.P. JSPS KAKENHI (17H06471) grant awarded to G.G. JDB laboratory was supported by FCT ERA-CAPS project EVOREPRO (ERA-CAPS-0001-2014). MBD laboratory is supported by ERC-2015-CoG-683258-Birth&Death. ET data acquisition was supported by a Cristian Boulin fellowship.

## Author contributions

S.G.P. designed and performed the experiments, together with A.L.S. (EM) and C.N. (sperm motility). T.P. analyzed SAS6, POC1 length distributions. S.G.P. and A.J.H. reconstructed and segmented the electron tomograms. M.S. supported ET data acquisition and tomogram reconstruction. G.G. provided the POC1-Citrine plasmid, preliminary plant genotypes and discussed experimental design. E.M.T. supervised the EM work and aided in ET data acquisition. S.G.P., J.D.B. and M.B.D. designed the study. S.G.P. wrote the first draft. S.G.P., J.D.B. and M.B.D. edited the manuscript with input from all authors. J.D.B. and M.B.D. supervised the work.

## Competing interests

The authors declare no competing interests.

## Supplementary Figures

**Figure S1.**
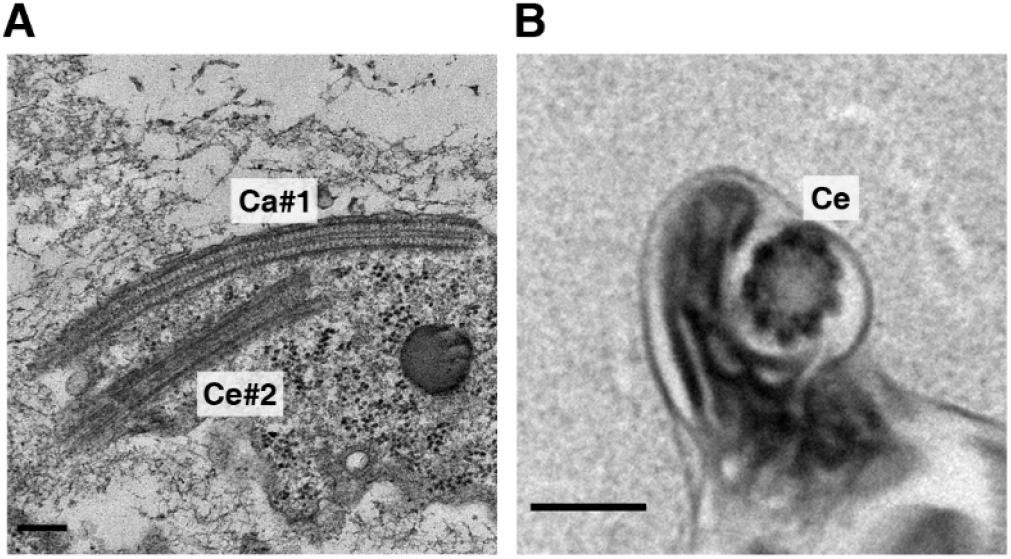
TEM images of particular features of developing sperm cells. **A,** Long cartwheel (Ca#1) region deprived of microtubules (“naked” cartwheel), while the other centriole (Ce#2) is anchored to the membrane and lacks a visible naked cartwheel. **B,** Thick section (300nm) showing a centriole (Ce) upon cell cytodifferentiation, without any clear cartwheel density in its lumen. Scale-bars = 200nm.

**Figure S2.**
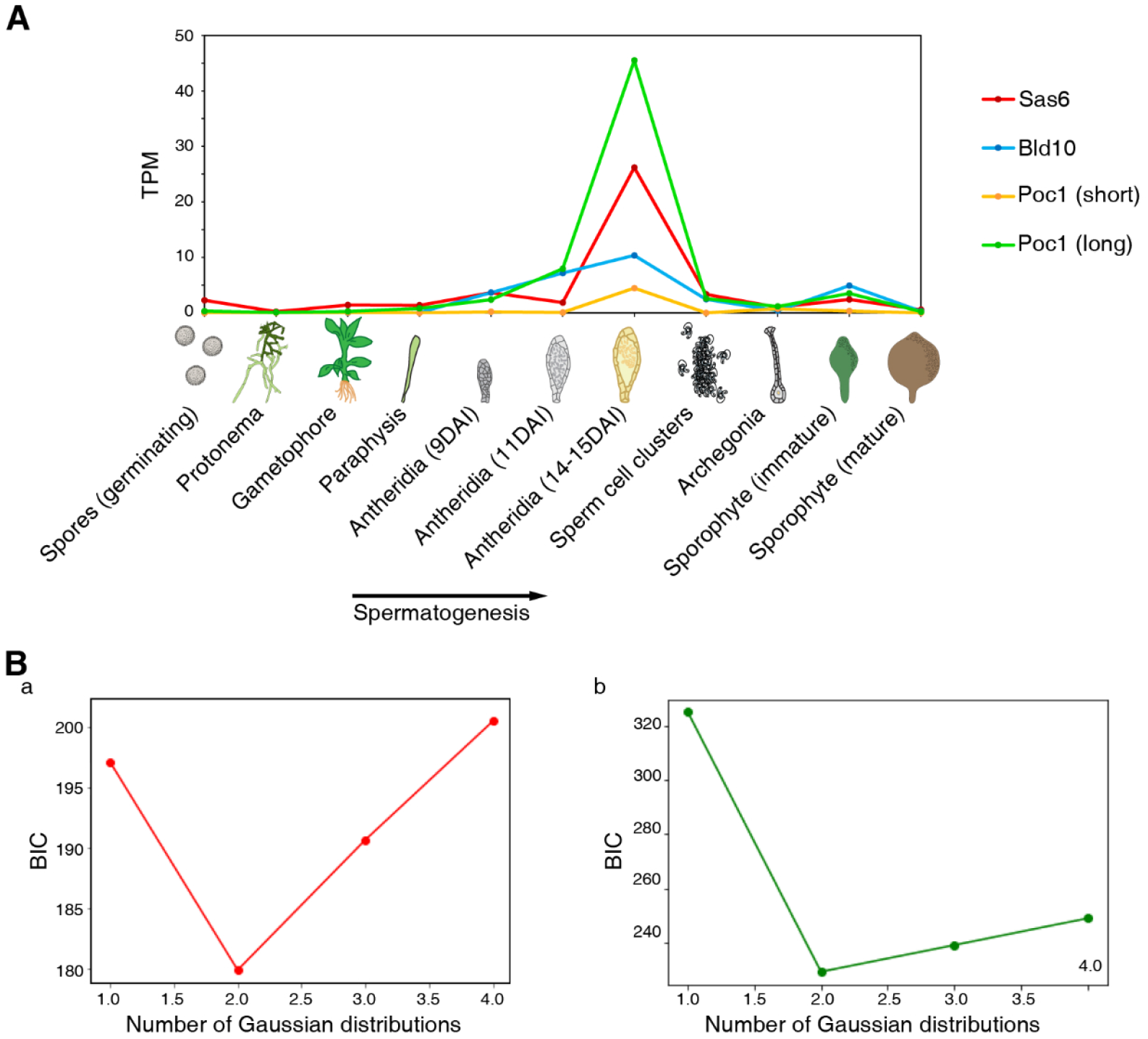
**A,** Gene expression profile of the conserved centriolar proteins SAS6 (Pp3c23_8210v3.1, red), BLD10 (Pp3c9_9030v3.1, blue), and POC1 (long transcript: Pp3c16_11590V3.1 - green; and shorter transcript: Pp3c16_11580V3.1 - yellow). Data collected from www.evorepro.plant.tools (Julca et al..). **B,** Bayesian information criterion (BIC) of Gaussian fitting to SAS6-mCherry (**Ba**) and POC1-Citrine (**Bb**) length distributions shown in Fig. 4D. Note that two is the number of Gaussian curves that result in the lowest BIC therefore supporting the existence of two distinct length populations.

**Figure S3.**
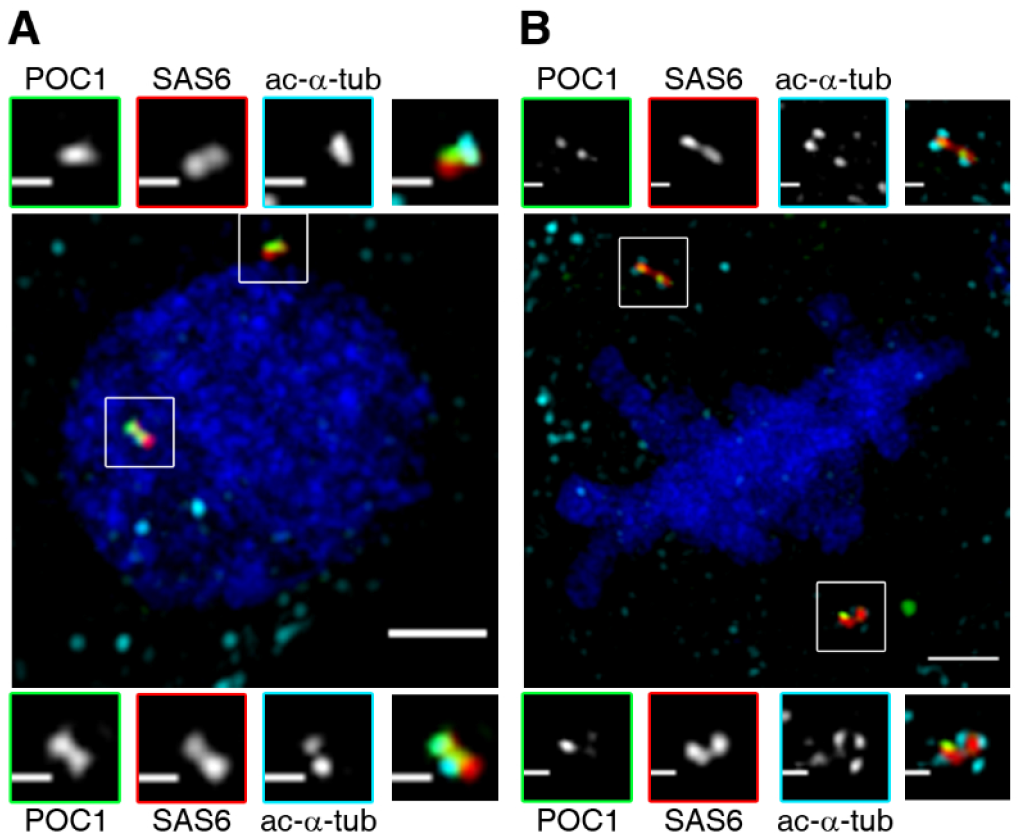
3D-SIM image showing the existence of two bicentrioles during interphase (**A**) and at the spindle poles during mitosis (**B**). Green – POC1-Citrine; Red – SAS6-mCherry; Cyan – acetylated-α-tubulin; Blue – DAPI. Scale-bars = 1μm or 250nm (insets).

**Figure S4.**
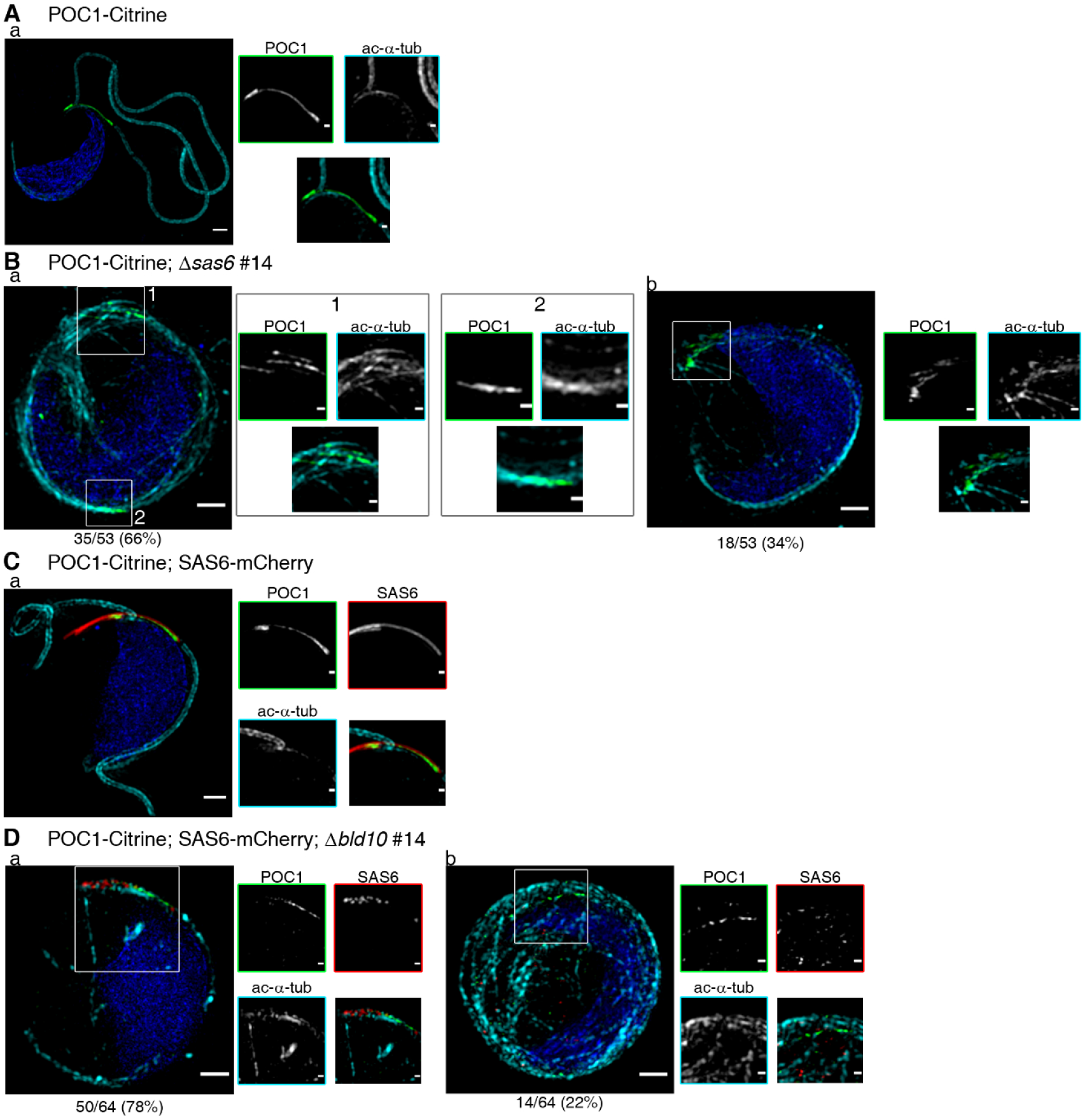
3D-SIM of analysis of stage IV sperm cells of the following genotypes: POC1-Citrine (**A,** genetic background for *Δsas6*), POC1-Citrine; *Δsas6* (**B**), POC1-Citrine; SAS6-mCherry (**C**, genetic background for *Δbld10*) and POC1-Citrine; SAS6-mCherry; *Δbld10* (**D**). **Aa,** POC1-Citrine plants show clear ciliary staining and centriole length asymmetry reported by POC1-Citrine signal. **Ba**, 66% of *Δsas6* stage IV. spermatids have two or more POC1 *foci*. **Bb,** In the remaining 34% of *Δsas6* cells, POC1 is concentrated in one tip of the cell but shows an abnormal organization. **Ca,** Stage IV POC1-Citrine; SAS6-mCherry, also showing clear ciliary staining, cartwheel elongation (given by SAS6-mCherry) and centriole asymmetry (reported by POC1-Citrine). **Da,** In the *Δbld10* sperm cells where SAS6 signal is observable (78%), POC1 localizes in the end of the SAS6 signal however, both signals are discontinuous. **Db,** When no clear SAS6 signal can be observed (22%), POC1 signal is found more widespread and discontinuous. Blue – DAPI; Cyan -acetylated-α-tubulin; Green – POC1-Citrine; Red – SAS6-mCherry (C and D). Scale-bars = 1μm or 250 nm in insets. For sample size considerations, see Table S1.

**Figure S5.**
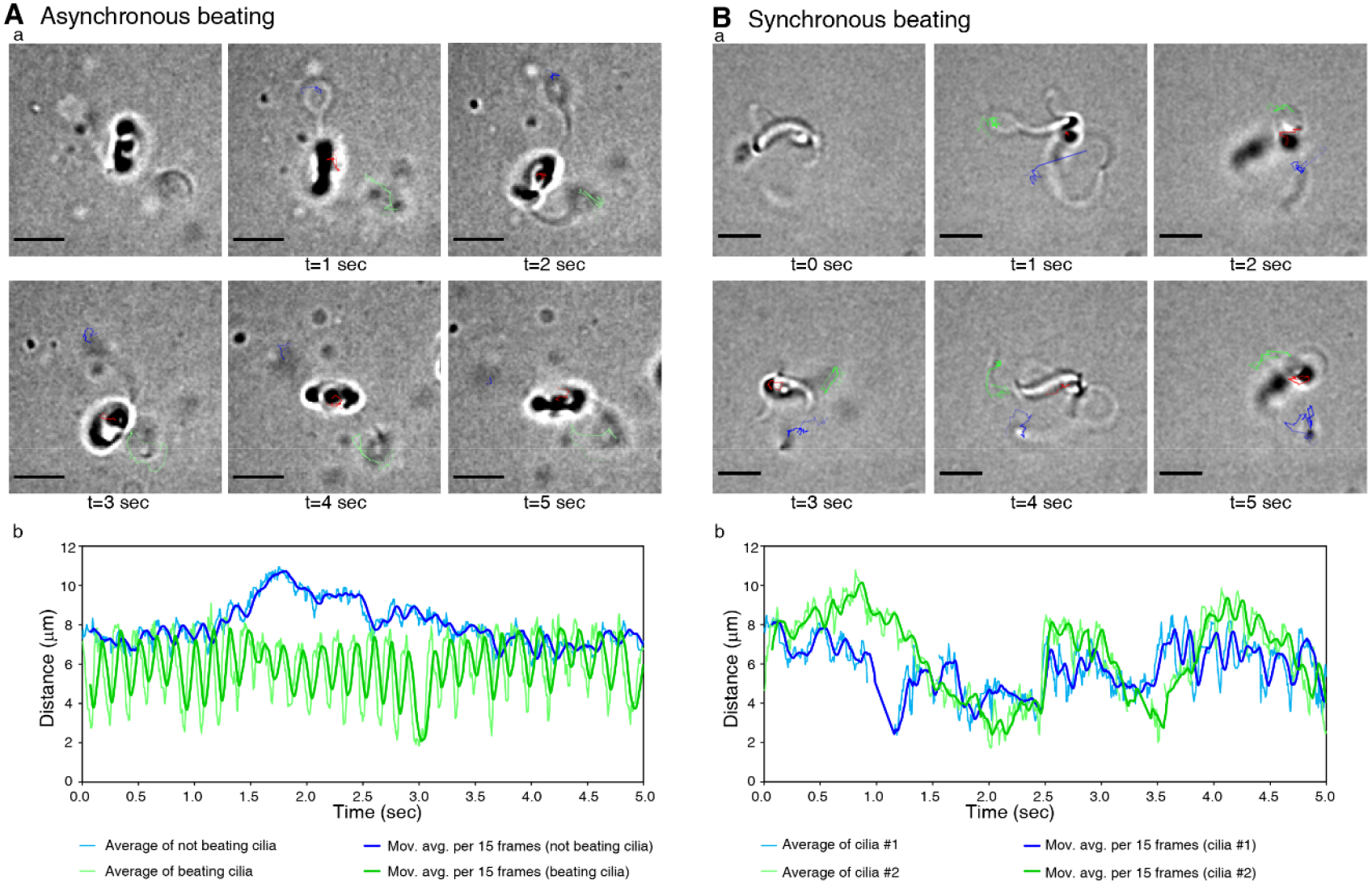
Ciliary beating in individual sperm cells. **A,** Sperm cell with only one active cilium (asynchronous beating, see movie 5). **Aa,** Snapshots of cilia and nuclear tracking over time. Motile cilium track is represented by the green line, inactive cilium by the blue and nucleus by the red line. Scale-bars = 5μm. **Ab,** Distance of ciliary tip of each cilium to the nucleus. Thin lines represent distance for the active (green) or inactive cilium (blue) while thicker lines represent the running averages for distances across 15 frames. **B,** Sperm cell with both cilia actively beating (synchronous beating, see movie 6). **Ba,** Snapshots of a cell tracked over time. Nuclear movement is represented by the green line, cilia are reported by red and blue lines, attributed randomly. Scale-bars = 5μm. **Bb,** Distance of each ciliary tip to the nucleus. Thin lines represent cilia distance while thicker lines represent its running averages across 15 frames.

## Supplementary movies and tables

**Movie 1** – Electron tomogram and respective 3D model of a bicentriole (Fig. 2A).

**Movie 2** – Electron tomogram and its 3D model of sister centrioles anchored to the MLS (Fig. 2B).

**Movie 3** – Electron tomogram and 3D model of asymmetrical sister centrioles anchored to the MLS and to the membrane during ciliogenesis (Fig. 3).

**Movie 4** – Qualitative assessment of sperm cell motility in control and K.O. plants (Fig. 5 and 6).

**Movie 5** – Example of an individual sperm cell displaying asynchronous ciliary beating (Fig. 7A and S5A).

**Movie 6** – Example of a sperm cell with both cilia actively beating synchronously (Fig. 7B and S5B).

**Table S1.** Plant genotypes used in this work and sample sizes considered.

**Table S2.** Primers, plasmids and plant genotypes used in this work.

## Notes

### Competing Interest Statement

The authors have declared no competing interest.

## References

Ashton, N.W., and Cove, D.J. (1977). The isolation and preliminary characterisation of auxotrophic and analogue resistant mutants of the moss, *Physcomitrella patens*. MGG Mol. Gen. Genet. 154, 87–95.

Aydogan, M.G., Wainman, A., Saurya, S., Steinacker, T.L., Caballe, A., Novak, Z.A., Baumbach, J., Muschalik, N., and Raff, J.W. (2018). A homeostatic clock sets daughter centriole size in flies. J. Cell Biol. 217, 1233–1248.

Bettencourt-Dias, M., Rodrigues-Martins, A., Carpenter, L., Riparbelli, M., Lehmann, L., Gatt, M.K., Carmo, N., Balloux, F., Callaini, G., and Glover, D.M. (2005). SAK/PLK4 is required for centriole duplication and flagella development. Curr. Biol. 15, 2199–2207.

Carothers, Z.B., and Duckett, J.G. (1980). The Bryophyte Spermatozoid : A Source of New Phylogenetic Information. Bull. Torrey Bot. Club 107, 281–297.

Carothers, Z.B., and Kreitner, G.L. (1968). Studies of Spermatogenesis in the Hepaticae. II. Blepharoplast structure in the spermatid of *Marchantia*. J. Cell Biol. 36, 603–616.

Carvalho-Santos, Z., Machado, P., Branco, P., Tavares-Cadete, F., Rodrigues-Martins, A., Pereira-Leal, J.B., and Bettencourt-Dias, M. (2010). Stepwise evolution of the centriole-assembly pathway. J. Cell Sci. 123, 1414–1426.

Gifford, E.M., and Larson, S. (1980). Developmental Features of the Spermatogenous Cell in *Ginkgo biloba*. Am. J. Bot. 67, 119–124.

Gopalakrishnan, J., Guichard, P., Smith, A.H., Schwarz, H., Agard, D.A., Marco, S., and Avidor-Reiss, T. (2010). Self-assembling SAS-6 multimer is a core centriole building block. J. Biol. Chem. 285, 8759–8770.

Le Guennec, M., Klena, N., Gambarotto, D., Laporte, M.H., Tassin, A.M., van den Hoek, H., Erdmann, P.S., Schaffer, M., Kovacik, L., Borgers, S., et al. (2020). A helical inner scaffold provides a structural basis for centriole cohesion. Sci. Adv. 6, eaaz4137.

Guichard, P., Chrétien, D., Marco, S., and Tassin, A.-M. (2010). Procentriole assembly revealed by cryo-electron tomography. EMBO J. 29, 1565–1572.

Guichard, P., Hamel, V., Le Guennec, M., Banterle, N., Iacovache, I., Nemcíková, V., Flückiger, I., Goldie, K.N., Stahlberg, H., Lévy, D., et al. (2017). Cell-free reconstitution reveals centriole cartwheel assembly mechanisms. Nat. Commun. 8, 14813.

Gupta, A., and Kitagawa, D. (2018). Ultrastructural diversity between centrioles of eukaryotes. J. Biochem. 164, 1–8.

Habedanck, R., Stierhof, Y.D., Wilkinson, C.J., and Nigg, E.A. (2005). The Polo kinase Plk4 functions in centriole duplication. Nat. Cell Biol. 7, 1140–1146.

Hilbert, M., Noga, A., Frey, D., Hamel, V., Guichard, P., Kraatz, S.H.W., Pfreundschuh, M., Hosner, S., Flückiger, I., Jaussi, R., et al. (2016). SAS-6 engineering reveals interdependence between cartwheel and microtubules in determining centriole architecture. Nat. Cell Biol. 18, 393–403.

Hiraki, M., Nakazawa, Y., Kamiya, R., and Hirono, M. (2007). Bld10p Constitutes the Cartwheel-Spoke Tip and Stabilizes the 9-Fold Symmetry of the Centriole. Curr. Biol. 17, 1778–1783.

Hodges, M.E., Scheumann, N., Wickstead, B., Langdale, J.A., and Gull, K. (2010). Reconstructing the evolutionary history of the centriole from protein components. J. Cell Sci. 123, 1407–1413.

Jerka-Dziadosz, M., Gogendeau, D., Klotz, C., Cohen, J., Beisson, J., and Koll, F. (2010). Basal body duplication in *Paramecium*: The key role of Bld10 in assembly and stability of the Cartwheel. Cytoskeleton 67, 161–171.

Joukov, V., and De Nicolo, A. (2019). The Centrosome and the Primary Cilium: The Yin and Yang of a Hybrid Organelle. Cells 8.

Julca, I., Ferrari, C., Flores-Tornero, M., Proost, S., Lindner, A.-C., Hackenberg, D., Steinbachová, L., Michaelidis, C., Gomes Pereira, S., Shekhar Misra, C., et al. Comparative transcriptomic analysis reveals conserved transcriptional programs underpinning organogenesis and reproduction in land plants. [BioRxiv preprint] DOI: 10.1101/2020.10.29.361501v1

Keller, L.C., Geimer, S., Romijn, E., John Yates, I., Zamora, I., and Marshall, W.F. (2009). Molecular Architecture of the Centriole Proteome: The Conserved WD40 Domain Protein POC1 Is Required for Centriole Duplication and Length Control. Mol. Biol. Cell 20, 1150–1166.

Khan, M.A., Rupp, V.M., Orpinell, M., Hussain, M.S., Altmüller, J., Steinmetz, M.O., Enzinger, C., Thiele, H., Höhne, W., Nürnberg, G., et al. (2014). A missense mutation in the PISA domain of HsSAS-6 causes autosomal recessive primary microcephaly in a large consanguineous pakistani family. Hum. Mol. Genet. 23, 5940–5949.

Khodjakov, A., Rieder, C.L., Sluder, G., Cassels, G., Sibon, O., and Wang, C.L. (2002). *De novo* formation of centrosomes in vertebrate cells arrested during S phase. J. Cell Biol. 158, 1171–1181.

Klena, N., Le Guennec, M., Tassin, A., van den Hoek, H., Erdmann, P.S., Schaffer, M., Geimer, S., Aeschlimann, G., Kovacik, L., Sadian, Y., et al. (2020). Architecture of the centriole cartwheel containing region revealed by cryo electron tomography. EMBO J. e106246.

Kleylein-Sohn, J., Westendorf, J., Le Clech, M., Habedanck, R., Stierhof, Y.D., and Nigg, E.A. (2007). Plk4-Induced Centriole Biogenesis in Human Cells. Dev. Cell 13, 190–202.

Kremer, J.R., Mastronarde, D.N., and McIntosh, J.R. (1996). Computer Visualization of Three-Dimensional Image Data Using IMOD. J. Struct. Biol. 116, 71–76.

Landberg, K., Pederson, E.R. a, Viaene, T., Bozorg, B., Friml, J., Jönsson, H., Thelander, M., and Sundberg, E. (2013). The moss *Physcomitrella patens* reproductive organ development is highly organized, affected by the two SHI/STY genes and by the level of active auxin in the SHI/STY expression domain. Plant Physiol. 162, 1406–1419.

LeGuennec, M., Klena, N., Aeschlimann, G., Hamel, V., and Guichard, P. (2020). Overview of the centriole architecture. Curr. Opin. Struct. Biol. 66, 58–65.

Levine, M.S., Bakker, B., Boeckx, B., Moyett, J., Lu, J., Vitre, B., Spierings, D.C., Lansdorp, P.M., Cleveland, D.W., Lambrechts, D., et al. (2017). Centrosome Amplification Is Sufficient to Promote Spontaneous Tumorigenesis in Mammals. Dev. Cell 40, 313–322.

Li, S., Fernandez, J.J., Marshall, W.F., and Agard, D.A. (2019). Electron cryo-tomography provides insight into procentriole architecture and assembly mechanism.

Lopes, C.A.M., Mesquita, M., Cunha, A.I., Cardoso, J., Carapeta, S., Laranjeira, C., Pinto, A.E., Pereira-Leal, J.B., Dias-Pereira, A., Bettencourt-Dias, M., et al. (2018). Centrosome amplification arises before neoplasia and increases upon p53 loss in tumorigenesis. J. Cell Biol. 217, 2353–2363.

Marteil, G., Guerrero, A., Vieira, A.F., De Almeida, B.P., Machado, P., Mendonça, S., Mesquita, M., Villarreal, B., Fonseca, I., Francia, M.E., et al. (2018). Over-elongation of centrioles in cancer promotes centriole amplification and chromosome missegregation. Nat. Commun. 9.

Mastronarde, D.N. (2005). Automated electron microscope tomography using robust prediction of specimen movements. J. Struct. Biol. 152, 36–51.

Mastronarde, D.N., and Held, S.R. (2017). Automated tilt series alignment and tomographic reconstruction in IMOD. J. Struct. Biol. 197, 102–113.

Matsuura, K., Lefebvre, P.A., Kamiya, R., and Hirono, M. (2004). Bld10p, a novel protein essential for basal body assembly in *Chlamydomonas*: Localization to the cartwheel, the first ninefold symmetrical structure appearing during assembly. J. Cell Biol. 165, 663–671.

Meehl, J.B., Bayless, B.A., Giddings, T.H., Pearson, C.G., and Winey, M. (2016). *Tetrahymena* Poc1 ensures proper intertriplet microtubule linkages to maintain basal body integrity. Mol. Biol. Cell 27, 2394–2403.

Mercey, O., Levine, M.S., LoMastro, G.M., Rostaing, P., Brotslaw, E., Gomez, V., Kumar, A., Spassky, N., Mitchell, B.J., Meunier, A., et al. (2019). Massive centriole production can occur in the absence of deuterosomes in multiciliated cells. Nat. Cell Biol. 21, 1544–1552.

Miki-Noumura, T. (1977). Studies on the *de novo* formation of centrioles: aster formation in the activated eggs of sea urchin. J. Cell Sci. 24, 203–216.

Miyamura, S., Matsunaga, S., and Hori, T. (2002). High-speed video microscopical analysis of the flagellar movement of *Marchantia polymorpha* sperm. Bryol. Res. 8, 79–83.

Moser, J.W., and Kreitner, G.L. (1970). Centrosome Structure in *Anthoceros laevis* and *Marchantia polymorpha*. J. Cell Biol. 44, 454–458.

Nabais, C., Pereira, S.G., and Bettencourt-Dias, M. (2017). Noncanonical Biogenesis of Centrioles and Basal Bodies. Cold Spring Harb. Symp. Quant. Biol. LXXXII.

Nabais, C., Pessoa, D., de-Carvalho, J., van Zanten, T., Duarte, P., Mayor, S., Carneiro, J., Telley, I., and Bettencourt-Dias, M. (2020). Plk4 triggers autonomous *de novo* centriole biogenesis and maturation. [BioRxiv preprint] DOI: 10.1101/2020.04.29.068650v2

Nakaoka, Y., Miki, T., Fujioka, R., Uehara, R., Tomioka, A., Obuse, C., Kubo, M., Hiwatashi, Y., and Goshima, G. (2012). An inducible RNA interference system in *Physcomitrella patens* reveals a dominant role of augmin in phragmoplast microtubule generation. Plant Cell 24, 1478–1493.

Nakazawa, Y., Hiraki, M., Kamiya, R., and Hirono, M. (2007). SAS-6 is a Cartwheel Protein that Establishes the 9-Fold Symmetry of the Centriole. Curr. Biol. 17, 2169–2174.

Nigg, E.A., and Holland, A.J. (2018). Once and only once: Mechanisms of centriole duplication and their deregulation in diseases. Nat. Rev. Mol. Cell Biol. 19, 297–312.

O’Toole, E.T., Giddings, T.H., McIntosh, J.R., and Dutcher, S.K. (2003). Three-dimensional Organization of Basal Bodies from Wild-Type and d-Tubulin Deletion Strains of *Chlamydomonas reinhardtii*. Mol. Biol. Cell 14, 2999–3012.

Ortiz-Ramírez, C., Michard, E., Simon, A.A., Damineli, D.S.C., Hernández-Coronado, M., Becker, J.D., and Feijó, J.A. (2017). GLUTAMATE RECEPTOR-LIKE channels are essential for chemotaxis and reproduction in mosses. Nature 549, 91–95.

Pearson, C.G., Osborn, D.P.S., Giddings, T.H., Beales, P.L., and Winey, M. (2009). Basal body stability and ciliogenesis requires the conserved component Poc1. J. Cell Biol. 187, 905–920.

Peel, N., Stevens, N.R., Basto, R., and Raff, J.W. (2007). Overexpressing Centriole-Replication Proteins *In Vivo* Induces Centriole Overduplication and *De Novo* Formation. Curr. Biol. 17, 834–843.

Pimenta-Marques, A., and Bettencourt-Dias, M. (2020). Pericentriolar material. Curr. Biol. 30, R687–R689.

Polin, M., Tuval, I., Drescher, K., Gollub, J.P., and Goldstein, R.E. (2009). *Chlamydomonas* swims with two “gears” in a eukaryotic version of run-and-tumble locomotion. Science (80). 325, 487–490.

Prigge, M.J., and Bezanilla, M. (2010). Evolutionary crossroads in developmental biology: *Physcomitrella patens*. Development 137, 3535–3543.

Rensing, S. a, Lang, D., Zimmer, A.D., Terry, A., Salamov, A., Shapiro, H., Nishiyama, T., Perroud, P.-F., Lindquist, E. a, Kamisugi, Y., et al. (2008). The *Physcomitrella* genome reveals evolutionary insights into the conquest of land by plants. Science (80). 319, 64–69.

Rensing, S.A., Goffinet, B., Meyberg, R., Wu, S.Z., and Bezanilla, M. (2020). The moss *Physcomitrium (Physcomitrella) patens*: A model organism for non-seed plants. Plant Cell 32, 1361–1376.

Renzaglia, K.S., and Garbary, D.J. (2001). Motile Gametes of Land Plants: Diversity, Development, and Evolution. CRC. Crit. Rev. Plant Sci. 20, 107–213.

Robbins, R.R. (1984). Origin and behavior of bicentriolar centrosomes in the bryophyte *Riella americana*. Protoplasma 121, 114–119.

Schorb, M., Haberbosch, I., Hagen, W.J.H., Schwab, Y., and Mastronarde, D.N. (2019). Software tools for automated transmission electron microscopy. Nat. Methods 16, 471–477.

Shaheen, R., Faqeih, E., Shamseldin, H.E., Noche, R.R., Sunker, A., Alshammari, M.J., Al-Sheddi, T., Adly, N., Al-Dosari, M.S., Megason, S.G., et al. (2012). POC1A truncation mutation causes a ciliopathy in humans characterized by primordial dwarfism. Am. J. Hum. Genet. 91, 330–336.

Shimamura, M., Brown, R.C., Lemmon, B.E., Akashi, T., Mizuno, K., Nishihara, N., Tomizawa, K.-I., Yoshimoto, K., Deguchi, H., Hosoya, H., et al. (2004). y - Tubulin in Basal Land Plants: Characterization, Localization, and Implication in the Evolution of Acentriolar Microtubule Organizing Centers. Plant Cell 16, 45–59.

Sorokin, S.P. (1968). Reconstructions of centriole formation and ciliogenesis in mammalian lungs. J. Cell Sci. 3, 207–230.

Stearns, T., Evans, L., and Kirschner, M. (1991). γ-Tubulin is a highly conserved component of the centrosome. Cell 65, 825–836.

Strnad, P., Leidel, S., Vinogradova, T., Euteneuer, U., Khodjakov, A., and Gönczy, P. (2007). Regulated HsSAS-6 Levels Ensure Formation of a Single Procentriole per Centriole during the Centrosome Duplication Cycle. Dev. Cell 13, 203–213.

Tinevez, J.Y., Perry, N., Schindelin, J., Hoopes, G.M., Reynolds, G.D., Laplantine, E., Bednarek, S.Y., Shorte, S.L., and Eliceiri, K.W. (2016). TrackMate: An open and extensible platform for single-particle tracking. Methods 115, 80–90.

Turner, F.R. (1970). The effects of colchicine on spermatogenesis in *Nitella*. J. Cell Biol. 46, 220–234.

Vaughn, K.C., Sherman, T.D., and Renzaglia, K.S. (1993). A centrin homologue is a component of the multilayered structure in bryophytes and pteridophytes. Protoplasma 175, 58–66.

Venoux, M., Tait, X., Hames, R.S., Straatman, K.R., Woodland, H.R., and Fry, A.M. (2012). Poc1A and Poc1B act together in human cells to ensure centriole integrity. J. Cell Sci. 126, 163–175.

