## Supplemental Table 1 for "The 3D architecture and molecular foundations of *de novo* centriole assembly *via* bicentrioles"

| Genotype | Line number | Electron microscopy (EM) | | | | | | | | | | | | | 3D Structured Illumination Microscopy (3D-SIM) | | | | Correlative Light and Electron Microscopy (CLEM) | Fertility rates | Sperm cell motility | | Main Figure |
|---|---|---|---|---|---|---|---|---|---|---|---|---|---|---|---|---|---|---|---|---|---|---|---|
| | | Chemically fixed<br>Cells/Structures, Blocks, Experiments | | | | | High pressure freezing + Automated<br>Cells/Structures, Blocks, Experiments | | | | | Electron Tomography (eT)<br>Cells/Structures, Blocks, Experiments | | | Cells, Experiments | | | | Cells/Structures, Blocks, Experiments | Independent counts | Motile vs. immotile<br>Clusters, Experiments | Ciliary beating tacking<br>Cells acquired, classified, tracked | |
| | | I | II | III | IV | V | I | II | III | IV | V | II | III | IV | I-II | III | IV | V | | | | | |
| WT Grandsden | | 10, 6, 3 | 3, 2, 2 | 76, 19, 5 | 148, 18, 4 | 120, 10, 3 | 2, 2, 2 | 1, 1, 1 | 40, 10, 4 | 54, 7, 3 | 23, 3, 2 | 3, 2, 1 | 3, 2, 1 | 3, 2, 1 | | | | | | 10 | 5, 3 | 135, 89, 10 | 1B, 2, 3, 7 |
| SAS6-mCherry; POC1-Citrine | 55 | | | | 21, 4, 2 | | | | | | | | | | 29, 5 | 16, 4 | 55, 4 | 38, 4 | | 25 | 3, 3 | | 4B, 4C |
| | 71 | | | | 9, 2, 1 | | | | | | | | | | 6, 2 | 8, 2 | 19, 2 | 8, 2 | | 5 | 7, 3 | | 4C |
| SAS6-mCherry | 304 | | | | | | | | | | | | | | 11, 6 | 8, 4 | 23, 6 | 14, 5 | 2, 2, 2 | 6 | | | 4C, 4D |
| γ-tubulin2-Citrine | 109 | | | | 53, 6, 2 | | | | | | | | | | 40, 11 | 19, 9 | 76, 13 | 16, 5 | | 20 | 7, 3 | | 5A, 5C |
| γ-tubulin2-Citrine; Δsas6 | 13 | | | | 70, 11, 3 | | | | | | | | | | 11, 5 | 14, 5 | 37, 4 | 13, 3 | | 5 | 5, 3 | | 5B, 5C |
| | 14 | | | | 15, 2, 1 | | | | | | | | | | 15, 4 | 11, 4 | 27, 3 | 16, 1 | | 5 | 6, 3 | | 5C |
| SAS6-mCherry; γ-tubulin2-Citrine | 78 | | | | 43, 7, 2 | | | | | | | | | | 54, 11 | 16, 7 | 94, 13 | 37, 4 | | 20 | 5, 3 | | 4C, 6A, 6D |
| | 61 | | | | | | | | | | | | | | 15, 6 | 10, 3 | 35, 6 | 9, 2 | | 5 | 4, 3 | | 4C |
| SAS6-mCherry; γ-tubulin2-Citrine; Δbld10 | 122 | | | | 70, 11, 3 | | | | | | | | | | 17, 6 | 20, 6 | 52, 6 | 21, 3 | | 5 | 4, 3 | | 6B, 6D |
| | 24 | | | | 24, 2, 1 | | | | | | | | | | 12, 3 | 17, 3 | 11, 2 | 7, 2 | | 5 | 4, 3 | | 6D |
| SAS6-mCherry; γ-tubulin2-Citrine; Δpoc1 | 10 | | | | 57, 7, 3 | | | | | | | | | | 18, 4 | 16, 4 | 30, 4 | 13, 4 | | 5 | 5, 4 | | 6C, 6D |
| | 21 | | | | 9, 2, 1 | | | | | | | | | | 15, 2 | 26, 2 | 23, 1 | 11, 1 | | 5 | 7, 3 | | 6D |
| POC1-Citrine | 341 | | | | 12, 3, 1 | | | | | | | | | | 13, 5 | 11, 4 | 62, 6 | 22, 4 | | 23 | 6, 3 | | 4C |
| POC1-Citrine; Δsas6 | 14 | | | | 18, 4, 1 | | | | | | | | | | 8, 4 | 20, 3 | 53, 4 | 24, 3 | | 7 | 8, 3 | | |
| | 70 | | | | 24, 3, 1 | | | | | | | | | | 5, 2 | 4, 2 | 16, 2 | 10, 2 | | 7 | 7, 3 | | *data not shown* |
| SAS6-mCherry; POC1-Citrine; Δbld10 | 14 | | | | 14, 2, 1 | | | | | | | | | | 25, 3 | 25, 3 | 64, 3 | 20, 3 | | 5 | 9, 3 | | |
| | 20 | | | | 10, 2, 1 | | | | | | | | | | 13, 1 | 15, 2 | 33, 2 | 13, 1 | | 5 | 7, 3 | | *data not shown* |

| **Suplementary Figure** |
|---|
| S1, S2A, S5 |
| S2B, S3, S4C |
| S2B, S3 |
| S2B |
| S2B |
| S2B |
| S2B, S4A |
| S4B |
| *data not shown* |
| S4D |
| *data not shown* |
