## Supplemental Table 2 for "The 3D architecture and molecular foundations of *de novo* centriole assembly *via* bicentrioles"

| Plant genotype | Reference | Genetic background for transformation | Plasmids used in transformation | HR plasmid<br>5' Arm | HR plasmid<br>Reporter protein/Resistence cassette | HR plasmid<br>3' Arm | gRNA sequences | Selection resistence | Genotyping<br>Mix 1 (gene/cassette integration) | Genotyping<br>Mix 2 (5' recombination) | Genotyping<br>Mix 3 (3' recombination) |
|---|---|---|---|---|---|---|---|---|---|---|---|
| **WT Grandsden** | #79 | | | | | | | | | | |
| **SAS6-mCherry** | This study | WT Grandsden | pBNRf_SAS6-mCherry_G418 | SAS6_GeneEnd_GA.For: AATTACCCTGTTATCCCTAGGCGTGCAAGCTGCTCAAAAGGTTC<br>SAS6_GeneEnd_GA.Rev: CAGCAGCACCAGCAGGGCTAGCCCTACGCGAGAAATGGCTCTTTGC<br>Ligated by Gibson Assembly | Linker-mCherry-NOSterminador; 35S::G418R<br>Ligated after endonuclease restriction (NheI + SpeI) in the Gibson Assembly step | SAS6_3END_GA_For: TGCTATACGAAGTTATACTAGTCTGTGCAAGAGCATTCGTTGTGAC<br>SAS6_3END_GA.Rev: CCCATGGATCGATGTTAACCGCCTATTGACCTAGCACAGTC<br>Ligated by Gibson Assembly | n/a | G418 | SAS6_genotypeF: GTGGTTTGGGAGTATTCAACACG<br>SAS6_genotypeR: CTAGGTTGGGGATGTTCTTCC | SAS6_genotypeF: GTGGTTTGGGAGTATTCAACACG<br>pBNR_Seq1_Rev: CACCTTGAAGCGCATGAACTC | 35Sfwd: GACGCACAATCCCACTATCC<br>SAS6_genotypeR: CTAGGTTGGGGATGTTCTTCC |
| **γ-tubulin2-Citrine** | #46 | | | | | | | | | | |
| **γ-tubulin2-Citrine; Δsas6** | This study | γ-tubulin2-Citrine | pBNR_SAS6_KO_Zeo_3UTR_c25A + pGENIOUS_Cas9_gRNA2_gRNA9 | pSAS6_KO_GAF: AAGCTAATTACCCTGTTATCCCTGCAGCATCTTTCTGTGTTCAGGTGC<br>pSAS6_KO_Zeo_GAR: GTTCGAACCCGGCTCTTTCCCTGATCAATC | Zeocin_pSAS6_GAF: GAGCCGGGTTCGAACGTACGTCGCG<br>Zeocin_3'SAS6_GAR: GAATGCTCTTGCACAGAGTTTAAACGCGTGGCGCCACTAGT<br>Ligated by Gibson Assembly, followed by site-directed mutagenesis (SAS6_3UTR_c25a_F: AGAGCATTCGTTGTGACACACGTTAAAGTGTCAGC) | SAS6_3END_GA_For: TGCTATACGAAGTTATACTAGTCTGTGCAAGAGCATTCGTTGTGAC<br>SAS6_3END_GA.Rev: cccatggatcgatgttaacCGCCTATTGACCTAGCACAGTC | gRNA#2: TAACGCTGACACTTTAACGTGGG<br>gRNA#9: TGATCAGGGAAAGAGCCGGATGG | Zeocin | pSAS6_genotype_F: GGCTGTATACTGCCACCTAAG<br>SAS6_genotypeR: CTAGGTTGGGGATGTTCTTCC | pSAS6_genotype_F: GGCTGTATACTGCCACCTAAG<br>35sZeo_rev: CGTCTTGATGAGACCTGCTG | SAS6_genotypeR: CTAGGTTGGGGATGTTCTTCC<br>SAS6_3end_c25a_F: GTTCCCTCACACCGGTGAC |
| **γ-tubulin2-Citrine; SAS6-mCherry** | This study | γ-tubulin2-Citrine | pBNR_SAS6-mCherry_HygR_3UTR_c25a + pGENIOUS_Cas9_gRNA2_gRNA1 | From pBNRf_SAS6-mCherry_G418 plasmid | Replaced G418R with HygR by endonuclease restriction with NheI+EcoRV from pBNRf_SAS6-mCherry_G418 plasmid<br>Followed by site directed mutagenesis (SAS6_3UTR_c25a_F: AGAGCATTCGTTGTGACACACGTTAAAGTGTCAGC) | From pBNRf_SAS6-mCherry_G418 plasmid | gRNA#1: TTTCTCGCGTAGGTAGCATGTGG<br>gRNA#2: TAACGCTGACACTTTAACGTGGG | Hygromycin B | SAS6_genotype_F: GTGGTTTGGGAGTATTCAACACG<br>SAS6_genotype_R: CTAGGTTGGGGATGTTCTTCC | SAS6_genotype_F: GTGGTTTGGGAGTATTCAACACG<br>pBNR_Seq1_Rev: CACCTTGAAGCGCATGAACTC | 35Sfwd: GACGCACAATCCCACTATCC<br>SAS6_genotype_R: CTAGGTTGGGGATGTTCTTCC |
| **γ-tubulin2-Citrine; SAS6-mCherry; Δbld10** | This study | γ-tubulin2-Citrine; SAS6-mCherry #78 | pBNR_BLD10_KO_Zeo_5end_g1057t + pGENIOUS_Cas9_gRNA4_gRNA3 | pBld10_KO_GAF: AAGCTAATTACCCTGTTATCCCTAGGGGGGTGAGGATTATGGAGG<br>pBld10_KO_Zeo_GAR: GTTCGAACCTGCAGGCCACCTAAGCTATGCCTG | Zeocin_pBld10_GAF: TAGGTGGCCTGCAGGTTCGAACGTACGTCGCG<br>Zeocin_3'Bld10_GAR: ATCTCTTAATTTCCATTCCGaCGCGTGGCGCCACTAGT<br>Ligated by Gibson Assembly, followed by site-directed mutagenesis (Bld10_KO_5end_g1057t_F: TATTATTCAGGCATAGCTTAGGTTGCCTGCAGGTTC) | BLD10_3END_GA.For: TGCTATACGAAGTTATACTAGTCGGAATGGAAATTAAGAGATCGAGAG<br>BLD10_3END_GA.Rev: CCCATGGATCGATGTTAACCAAAGAGGTAGCACAAGGATCAAGGTTCA | gRNA#3: TTATTCAGGCATAGCTTAGGTGG<br>gRNA#4: GAGTCCGAACTAAAAATGATCGG | Zeocin | pBLD10_genotype: CGGGAAGCTCTCTGTAGGATTG<br>Bld10_genotype_R: CAATCTTTGTGCCAGCTTCCTGC | pBLD10_genotype: CGGGAAGCTCTCTGTAGGATTG<br>35sZeo_rev: CGTCTTGATGAGACCTGCTG | SAS6_3end_c25a_F: GTTCCCTCACACCGGTGAC<br>Bld10_genotype_R: CAATCTTTGTGCCAGCTTCCTGC |
| **γ-tubulin2-Citrine; SAS6-mCherry; Δpoc1** | This study | γ-tubulin2-Citrine; SAS6-mCherry #78 | pBNR_POC1_KO_Zeo + pGENIOUS_Cas9_gRNA14_gRNA13 | PstI_POC1_5UTR_F: CTGCAGCACGCAAAAGCTAGAGCAAG<br>POC1_5UTR_XhoI_R: CTCGAGTGAGGAGTCCATAAAGTGAGTTGCC<br>Inserted after endonuclease restriction (PstI + XhoI) and TA ligation | 35S::BleoR<br>Ligated by TA ligation after endonuclease restriction (XhoI + MluI) | MluI_POC1_3UTR_F: ACGCGTTCGTGGCAATTGATGCGAAG<br>POC1_3UTR_PacI_R: TTAATTAACCATTCGGCAGTCGTTACTG<br>Inserted after endonuclease restriction (MluI + PacI) and TA ligation | gRNA#13: ATACGTCGTATATCTTTCCTGGG<br>gRNA#14: GGACTGCAGGGGATTAATCGTGG | Zeocin | pPOC1_genotype_F: CTAGTGCGTAACACATCCTGG<br>pPOC1_genotype_R: CTTTGTAGCGGGAGCTTGATTAG | pPOC1_genotype_F: CTAGTGCGTAACACATCCTGG<br>35sZeo_rev: CGTCTTGATGAGACCTGCTG | SAS6_3end_c25a_F: GTTCCCTCACACCGGTGAC<br>pPOC1_genotype_R: CTTTGTAGCGGGAGCTTGATTAG |
| **POC1-Citrine** | This study | WT Grandsden | EKv194(Pp3c16_11590(POC1)-CITRINE) + pGENIOUS_cas9_U6_gRNA14 | POC1_CTRN_IF_F: GAGGTCGACGGTATCGTACTTGTCGTTACTCCCAGG<br>POC1_CTRN_IF_R: CAAGATATCAAGCTTATCCCCTGCAGTCCTTGGTC | Citrine_F: AAGCTTGATATCTTGGTGAGCAA<br>Citrine_R: ATAGGGACTTTAGGAGATCTGGA<br>In Fusion cloning with backbone pCTRN-NPTII2 amplified using pCTRN-Npt II-2 F (ACGAGACGACTAAACCTGGAGCC) and pCTRN-Npt II-2 R (GATACCGTCGACCTCGAGGGGGG) | POC1 3'UTR CTRN IF F: TCCTAAAGTCCCTATTCGTGGCAATTGATGCGAAGC<br>POC1 3'UTR CTRN IF R: GTTTAGTCGTCTCGTCGCCTTCACAAGCCTGCACG | gRNA#14: GAGTCCGAACTAAAAATGATCGG | G418 | POC1GeneEnd_genotype_F: CGGCCACTCTCTAGAGAAGG<br>pPOC1_genotype_R: CTTTGTAGCGGGAGCTTGATTAG | POC1GeneEnd_genotype_F: CGGCCACTCTCTAGAGAAGG<br>mCardinal_nSTOP_R: CTTGTACAGCTCGTCCATGCC | 35Sfwd: GACGCACAATCCCACTATCC<br>pPOC1_genotype_R: CTTTGTAGCGGGAGCTTGATTAG |
| **POC1-Citrine; Δsas6** | This study | POC1-Citrine #341 | pBNR_SAS6_KO_Zeo_3UTR_c25A + pGENIOUS_Cas9_gRNA2_gRNA9 | Same as for γ-tubulin2-Citrine; Δsas6 | | | | | | | |
| **POC1-Citrine; SAS6-mCherry** | This study | POC1-Citrine #341 | pBNR_SAS6-mCherry_HygR_3UTR_c25a + pGENIOUS_Cas9_gRNA2_gRNA1 | Same as for γ-tubulin2-Citrine; SAS6-mCherry | | | | | | | |
| **POC1-Citrine; SAS6-mCherry; Δbld10** | This study | POC1-Citrine; SAS6-mCherry #55 | pBNR_BLD10_KO_Zeo_5end_g1057t + pGENIOUS_Cas9_gRNA4_gRNA3 | Same as for γ-tubulin2-Citrine; SAS6-mCherry; Δbld10 | | | | | | | |
